# Post-synaptic morphology of mouse neuromuscular junctions is linked to muscle fibre type

**DOI:** 10.1101/2020.04.04.025106

**Authors:** Aleksandra M. Mech, Anna-Leigh Brown, Giampietro Schiavo, James N. Sleigh

## Abstract

The neuromuscular junction (NMJ) is the highly specialised peripheral synapse formed between lower motor neuron terminals and muscle fibres. Post-synaptic acetylcholine receptors (AChRs), which are found in high density in the muscle membrane, bind to acetylcholine released into the synaptic cleft of the NMJ, ultimately facilitating the conversion of motor action potentials to muscle contractions. NMJs have been studied for many years as a general model for synapse formation, development and function, and are known to be early sites of pathological changes in many neuromuscular diseases. However, information is limited on the diversity of NMJs in different muscles, whether muscle fibre type impacts NMJ morphology and growth, and the relevance of these parameters to neuropathology. Here, this crucial gap was addressed using a robust and standardised semi-automated workflow called NMJ-morph to quantify features of pre- and post-synaptic NMJ architecture in an unbiased manner. Five wholemount muscles from wild-type mice were dissected and compared at immature (post-natal day, P7) and early adult (P31-32) timepoints. Post-synaptic AChR morphology was found to be more variable between muscles than that of the motor neuron terminal and there were greater differences in the developing NMJ than at the mature synapse. Post-synaptic architecture, but not neuronal morphology or post-natal synapse growth, correlates with fibre type and is largely independent of muscle fibre diameter. Counter to previous observations, this study indicates that smaller NMJs tend to innervate muscles with higher proportions of fast twitch fibres and that NMJ growth rate is not conserved across all muscles. Furthermore, healthy pre- and post-synaptic NMJ morphological parameters were collected for five anatomically and functionally distinct mouse muscles, generating reference data that will be useful for the future assessment of neuromuscular disease models.

**Graphical Abstract:** 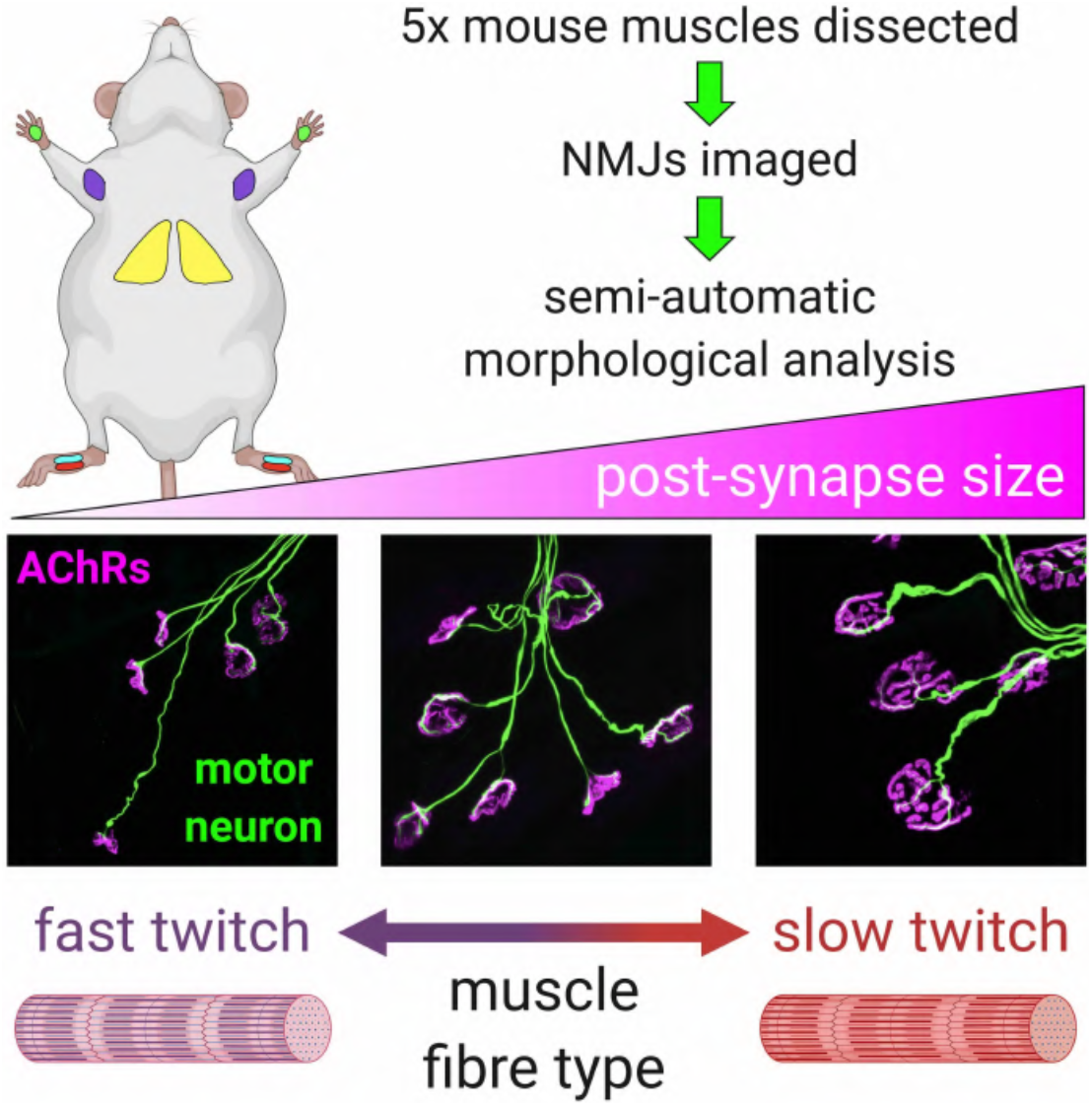

## 1. Introduction

The neuromuscular junction (NMJ) is a peripheral synapse formed between lower motor neurons and skeletal muscle fibres that has been studied in many different species including humans (Cramer and Van Essen, 1995; Campbell and Ganetzky, 2012; Grice *et al.*, 2018; Hammond *et al.*, 2016; Jones *et al.*, 2017; Kuno *et al.*, 1971; Nyström, 1968; Prakash *et al.*, 1996; Sakowski *et al.*, 2012). Upon action potential-mediated depolarisation of the pre-synaptic neuronal membrane, the neurotransmitter acetylcholine (ACh) is released into the NMJ synaptic cleft, where it binds to ACh receptors (AChRs) found in high density on post-synaptic muscle membranes in direct apposition to motor nerve terminals. This results in sodium ion influx into the muscle, post-synaptic membrane depolarisation and, if a threshold is reached, excitation-contraction coupling and muscle contraction. Due to its experimental accessibility and large size, the NMJ has been used for many years as a model for pre- and early post-natal synapse development (Sanes and Lichtman, 1999). During this time, the mammalian NMJ undergoes a series of growth and maturation processes that refine the synaptic architecture and enhance neurotransmission efficiency to facilitate motor performance (Shi *et al.*, 2012).

At birth, mammalian NMJs are polyinnervated, meaning that the endplate is contacted by more than one motor nerve ending, and in some instances more than ten different axons (Tapia *et al.*, 2012). During early post-natal development, equating to less than two weeks in mice, asynchronous axon withdrawal occurs and NMJs become monoinnervated. Known as synapse elimination, this process is thought to refine peripheral nerve architecture and is driven by activity, inter-nerve competition, and reciprocal nerve-muscle-glial signalling (Favero *et al.*, 2012; I. W. Smith *et al.*, 2013; Turney and Lichtman, 2012). This is accompanied by significant changes in the post-synapse to maintain the close proximity of motor terminals to AChRs. For example, the density and number of AChRs increase and the morphology of AChR clusters transforms from a simple, plaque-like shape to a complex, multi-perforated, pretzel-like structure (Marques *et al.*, 2000). This is, at least in part, due to remodelling of the post-synaptic membrane to create junctional folds, enlarging the contact area within the synaptic cleft. Moreover, the molecular composition of the membrane also changes to assure correct functioning of the mature synapse; for instance, the foetal AChR γ subunit is exchanged for the adult ε subunit (Missias *et al.*, 1996).

Mammalian muscles are composed of two main classes of force-generating (extrafusal) muscle fibre, ‘slow twitch’ (type I) and ‘fast twitch’ (type II), which have differing anatomical, physical and metabolic properties (Schiaffino and Reggiani, 2011). Slow twitch fibres generate energy through oxidative metabolism and can be continuously used for long periods of time (*i.e.* hours), therefore they function primarily to maintain body posture. Based on differential expression of myosin heavy chain (MHC) genes, fast twitch fibres can be further subdivided into three principal categories: type IIa (expressing MYH2), IIx (expressing MYH1) and IIb (expressing MYH4) (Talbot and Maves, 2016). The time to contraction and time to fatigue of fast twitch fibres is shorter than slow twitch, and they lie on a spectrum of decreasing times from IIa to IIb. Type IIa fibres are oxidative and most similar to slow twitch fibres, while type IIx and IIb are both glycolytic, with IIb fibres being required for the shortest and most intense movements. There are conflicting reports as to size categories of each fibre subtype, which is probably due to their diameters fluctuating between types of muscle; for example, slow twitch fibres are larger than fast twitch in the rat soleus muscle, which is composed of 86-93% slow twitch fibres, but smaller in the rat extensor digitorum longus (EDL) muscle that is made up of only 2-3% slow twitch (Alnaqeeb and Goldspink, 1987). Nevertheless, slow twitch fibres are more often reported as being smaller than fast twitch, with type IIb being the largest (Stifani, 2014). In addition to intrinsic disparities, muscle fibre types are innervated by distinct classes of alpha motor neuron (Stifani, 2014), the firing patterns of which appear to impact the contractile properties of fibres (Pette and Vrbová, 1985), while the fibre also appears to be able to retrogradely influence the identity of their innervating neuron (Chakkalakal *et al.*, 2010). The motor neurons that innervate slow twitch fibres generally have smaller cell bodies and axons, require a lower threshold to stimulation, and have slower conduction speeds, whereas fast-fatiguable motor neurons innervating type IIb fibres are at the other end of the spectra for these characteristics (Stifani, 2014).

The size of NMJs has been reported to correlate with fibre diameter in several different species and muscles (Balice-Gordon and Lichtman, 1990; Harris, 1954; Kuno *et al.*, 1971; Nyström, 1968). Moreover, manipulations of muscle fibre size can cause equivalent changes in the NMJ (Balice-Gordon *et al.*, 1990). However, this relationship between size of NMJs and fibres may only be observable when fibre classes are individually assessed (Prakash *et al.*, 1995a; Waerhaug and Lømo, 1994) or within certain types of muscle (Jones *et al.*, 2016). Indeed, similar to muscle fibre diameters, inconsistencies in morphological features of NMJs associated with the distinct fibre types have been reported. For instance, using both light and electron microscopy, the average area of NMJs in predominantly slow twitch soleus muscles of the rat was shown to be larger than in the fast twitch EDL (McArdle and Sansone, 1977; Wood and Slater, 1997). Contrasting with this, NMJs innervating type I fibres in rat diaphragm muscle were shown to be less than half the size of those innervating type II fibres (Prakash *et al.*, 1996). However, correcting for fibre diameter, the NMJ size does not vary quite so much across the fibre types and muscles, although type I fibre NMJs are larger than fast twitch NMJs in rat diaphragm muscle (Prakash *et al.*, 1996, 1995b; Prakash and Sieck, 1998). This suggests that features other than fibre diameter contribute to NMJ size differences between muscle fibre types, e.g. the innervating motor neuron and its firing properties (Waerhaug and Lømo, 1994).

The NMJ is an early and important site of pathology in many different neuromuscular conditions, including congenital myasthenic syndromes (Engel *et al.*, 2015), autoimmune diseases such as myasthenia gravis (Gilhus, 2012), and neurodegenerative disorders including amyotrophic lateral sclerosis (Williamson *et al.*, 2019), spinal muscular atrophy (Murray *et al.*, 2008), and Charcot-Marie-Tooth disease (Spaulding *et al.*, 2016). Furthermore, aging appears to result in structural alterations at the rodent NMJ (Gonzalez-Freire *et al.*, 2014), while axotomy causes NMJ dismantling prior to axonal degeneration (Miledi and Slater, 1970). Consequently, it is of considerable importance to be able to accurately assess the integrity of neuromuscular synapses in models of pathology. Some of the first published attempts to visualise the NMJ date back to the late 19th century and used gold and silver preparations to determine endplate shape (Kühne, 1887). Development of cholinesterase staining (Koelle and Friedenwald, 1949), followed by the advent of immunofluorescence techniques and confocal microscopy allowed for independent assessment of pre- and post-synaptic NMJ components. Pathological NMJ phenotypes (e.g. denervation counts and neurofilament accumulation) are often assessed by eye, possibly reducing objectivity and limiting inter-study comparisons. However, this issue has been tackled by the Gillingwater laboratory, who have developed a free, ImageJ-based methodology called NMJ-morph for comparative analysis of neuromuscular synapses (Jones *et al.*, 2016), which has been successfully used to evaluate NMJs in patients with cancer cachexia (Boehm *et al.*, 2020). By using thresholded, binary images, data on 20 NMJ morphological variables can be quickly and reliably obtained, with the proviso that several are not independent (e.g. nerve terminal perimeter and area). Operator-independent information on the pre-synaptic nerve terminal (e.g. number of branches and nerve area), post-synaptic AChRs (e.g. perimeter and area), and overlapping features (e.g. synaptic contact area) can be objectively generated for accurate cross-muscle, cross-study and even cross-laboratory comparisons.

Here, the same software was used to assess morphology of neuromuscular synapses at two different timepoints, representing a developing (postnatal day 7, P7) and a mature (P31-32) NMJ. Performing analyses in five different wholemount muscles with varying percentages of slow- and fast twitch fibres, this novel morphometric approach was used to assess the influence of fibre type and diameter on pre- and post-synaptic NMJ morphology, as well as post-natal synaptic development/growth.

## 2. Materials and Methods

### 2.1 Animals

Wild type mice on a C57BL/6J background were sacrificed at P7 and P31-32 for post-natal developmental and early adult timepoints, respectively. Muscles were harvested from three males and three females at each age, weighting an average of 4.0 g at P7 and 17.0 g at P31-32 (**Supplementary Table 1**). P7 animals were taken from a single litter, whereas P31-32 mice came from two different litters. All muscles were dissected from the left side of the body, except for lumbricals of the hand, which were taken from the right side. Mouse handling and experiments were performed under license from the UK Home Office in accordance with the Animals (Scientific Procedures) Act (1986) and approved by the University College London – Queen Square Institute of Neurology Ethics Committee.

### 2.2 Tissue preparation and immunohistochemistry

TVA, ETA, FDB, and lumbrical muscles of both the hands and feet were dissected (Murray *et al.*, 2014; Sleigh *et al.*, 2014a; Tarpey *et al.*, 2018) and stained as previously described (Sleigh *et al.*, 2017, 2014b). Briefly, muscles were excised and fixed for 10 min in 4% (w/v) paraformaldehyde (PFA, Electron Microscopy Sciences, Hatfield, PA) in phosphate buffered saline (PBS), before being permeabilised for 30 min in 2% (v/v) Triton X-100 in PBS and blocked for 30 min in 4% (w/v) bovine serum albumin and 1% (v/v) Triton X-100 in PBS. Muscles were then probed overnight at 4°C in blocking solution with 1/250 mouse anti-neurofilament (2H3, Developmental Studies Hybridoma Bank [DSHB], Iowa City, IA, supernatant) and 1/25 mouse pan anti-synaptic vesicle 2 (SV2, DSHB, supernatant) antibodies to co-visualise axons and pre-synaptic nerve terminals, respectively. The following day muscles were washed several times with PBS before being incubated with 1/1,000 Alexa Fluor 488 donkey anti-mouse IgG secondary antibody (Life Technologies, Carlsbad, CA, A-21202) and 1/1,000 Alexa Fluor 555 α-bungarotoxin (α-BTX, Life Technologies, B35451) in PBS for 2 h. Muscles were washed with PBS and then mounted in Fluoromount-G (Southern Biotech, Birmingham, AL, 0100-01) on microscope slides for imaging. All five muscles from each mouse were processed in parallel.

### 2.3 NMJ imaging

NMJs were imaged using an LSM 510 laser scanning microscope (Zeiss, Oberkochen, Germany) with Argon/2 and HeNe543 lasers for neurons (labelled green; 2H3/SV2) and post-synaptic AChRs (labelled red; α-BTX), respectively. Z-stacks with a 1 μm interval were taken using a Plan-Apochromat 63x/1.4 oil immersion objective. Images of P31-32 samples were taken with 0.7x zoom and images of P7 samples with 2x zoom.

### 2.4 NMJ analysis

ImageJ software (https://imagej.nih.gov/ij/) combined with the NMJ-morph workflow (Jones *et al.*, 2016) was used to describe 19 morphological variables of NMJs across all five muscles, each individually described below. The resolution of analysed images was 512 × 512 pixels, corresponding to 101.02 × 101.02 μm at P31-32 and 35.71 × 35.71 μm at P7. For all variables, 20 *en face* NMJs with clearly visible pre-synaptic axons and terminals were assessed (resulting in analysis of 1,200 NMJs in total). A twentieth variable of polyinnervation percentage was visually assessed at 100 NMJs per muscle (resulting in analysis of 6,000 NMJs in total). In the case of lumbrical and FDB muscles, where more than one muscle was processed, NMJs were assessed across all muscles (as were muscle fibres, see below). All analyses were performed on maximum intensity Z-stack projections and no samples were excluded from analyses once imaged.

#### 2.4.1 Initial image processing

Maximum intensity Z-stack projections were created using ImageJ (Image > Stacks > Z Project… > Projection type: Max Intensity). The two separate fluorophore channels were split (Image > Colour Split Channels) and a threshold selected (Image > Adjust > Threshold…). Of several available thresholding methods, three provided the most accurate binary representation of NMJs. ‘Huang’, ‘Yen’ or manual method were chosen based on how well they resembled the original image. The ‘Huang’ method was used for 56 and 79 percent of NMJs from P31-32 and P7, respectively, the ‘Yen’ method for 42 and 1 percent, and the manual method for 2 and 20 percent. Images were then cleaned from background (Process > Noise > Despeckle) and saved as binary images (Process > Binary > Make Binary) with green and red backgrounds for pre- and post-synaptic images, respectively. With the exception of ‘Number of AChR clusters’, all following analyses were performed on these ‘clean’ images.

#### 2.4.2 Pre-synaptic measurements

Polyinnervation percentage was calculated by determining the proportion of NMJs with more than one motor axon innervating the post-synaptic endplate. For NMJ-morph analyses, axons were manually erased with the paintbrush tool, leaving behind nerve terminals whose ‘Area’ and ‘Perimeter’ were taken (Edit > Selection > Create Selection > Analyze > Measure). The remaining nerve terminal was then turned into a mask (Process > Binary > Convert to Mask) and skeletonized (Process > Binary > Skeletonize) to allow for the use of the Binary Connectivity plugin (Plugins > Morphology > Binary Connectivity), which created a white binary image on the white background. Next, the image histogram and the list of values corresponding to the number of pixels on the x-axis of the histogram were opened (Analyse > Histogram > List). Histogram values 0, 2, 4 and 5, equating to the numbers of background white pixels, number of terminal pixels, number of three-point branch pixels and number of four-point branch pixels, respectively, were noted. This allowed for automatic calculation of ‘Number of terminal branches’, ‘Number of branch points’, ‘Total length of branches’, ‘Average length of branches’ and ‘Complexity’. ‘Complexity’ was determined by calculating log10[‘Number of terminal branches’ × ‘Number of branch points’ × ‘Total length of branches’]. ‘Axon diameter’ was measured using the line tool at the edge of endplates and at the points of maximum and minimum axon diameter (Analyze > Measure). The mean of three measurements was recorded.

#### 2.4.3 Post-synaptic measurements

For clarity, ‘endplates’, ‘AChRs’ and ‘AChR clusters’ are slightly different post-synaptic structures of decreasing size. Endplates include the post-synaptic staining, along with associated perforations and other areas devoid of α-BTX that occur within the perimeter of staining. Hence, endplate area is a good proxy for overall NMJ size. AChRs represent the footprint of post-synaptic staining alone (*i.e.* excluding α-BTX-negative areas). AChR clusters include only the α-BTX staining found in discrete groupings.

‘Clean’ post-synaptic images were opened. ‘Perimeter’ and ‘Area’ of AChR clusters were recorded (Edit > Selection > Create Selection > Analyze > Measure). To obtain the ‘Diameter’, ‘Perimeter’ and ‘Area’ of the endplate, images were processed to create the background (Process > Subtract Background… > Create background (rolling ball radius: 50.0 pixels) > Process > Binary > Make Binary). The line tool was used to draw the maximum ‘Endplate diameter’ (Analyze > Measure) and automatic outline tool to gather ‘Area’ and ‘Perimeter’ (Edit > Selection > Create Selection > Analyze > Measure). Values were noted and ‘Compactness’ calculated automatically [‘AChR Area’ divided by ‘Endplate Area’ × 100]. ‘Number of AChR clusters’, ‘Area of AChR clusters’ and ‘Fragmentation’ were calculated from maximum intensity projection images of post-synaptic terminals (Image > Stacks > Z Project… > Projection type: Max Intensity; then Image Colour > Split channels > Endplate channel; then Process > Binary > Make Binary; then Process > Find Maxima… > Segmented particles). The number of segmented particle clusters was manually counted and noted down, which allowed for calculation of ‘Area of AChR clusters’ and ‘Fragmentation’. Fragmentation was calculated with the following formula [1−(1/‘Number of AChR clusters’)].

#### 2.4.4 Combined pre- and post-synaptic measurements

To calculate ‘Area of synaptic contact’ and ‘Overlap’ between nerve terminals and endplates, both pre- and post-synaptic ‘clean’ images were opened and then merged, which enabled visualisation and thus recording of the ‘Area of unoccupied AChRs’ (For endplate: Edit > Invert; then Image > Stacks > Tools > Concatenate…; then Image > Stacks > Z Project… > Projection type: Average intensity; then Process > Binary > Make Binary; then Edit > Selection > Create Selection > Analyze > Measure). ‘Area of synaptic contact’ was calculated using the following formula [‘AChR Area’ – ‘Area of unoccupied AChRs’], and ‘Overlap’ with [(‘Area of synaptic contact’/‘AChR Area’) × 100].

### 2.5 Muscle fibre diameter analysis

After imaging P31-32 NMJs for NMJ-morph analysis, microscope slides were immersed in PBS to facilitate coverslip removal. Muscles were then teased using forceps to separate and aid identification of individual fibres, before re-mounting in Fluoromount-G on new microscope slides. Muscles were imaged at a resolution of 1024 × 1024 by differential interference contrast microscopy on an inverted LSM780 laser scanning microscope (Zeiss) with a 20x objective. Using ImageJ, fibre diameters were determined by taking five width measurements at different points along individual fibres. Damaged fibres were avoided, as were endings that were patently larger than the rest of the fibre. 30 fibres were measured per muscle per animal to generate mean values (resulting in analysis of 900 fibres in total). Given the method of processing, the analysed fibres were highly unlikely to be those in which NMJ morphology was assessed.

### 2.6 Statistical analysis

Data were assumed to be normally distributed unless evidence to the contrary could be provided by the D’Agostino and Pearson omnibus normality test. GraphPad Prism 8 (version 8.4.0, La Jolla, CA) was used for all statistical analyses. Means ± standard error of the mean are plotted in **Figures 2-4** and **Supplementary Figures 2-6**, as well as individual data points generated from each individual mouse (*i.e.* the mean values from 20 NMJs/30 fibres) in **Figure 2** and **3** and **Supplementary Figure 4B**, **6A** and **6C**. In all bar charts, muscles are plotted on the x-axes in ascending order of the percentage of fast twitch fibres (*i.e.* 68% in the TVA to 100% in the hand lumbricals, see **Table 1**). Moreover, the colour scheme for the five muscles is maintained throughout all figures and supplementary figures (yellow, TVA; cyan, foot lumbricals; purple, ETA; red, FDB; green, hand lumbricals). Individual morphological variables were statistically analysed using a repeated-measures one-way analysis of variance (ANOVA) test in **Figure 2** and **3** and **Supplementary Figure 4B**, **6A** and **6C**. This test was used because the five dissected muscles were all taken from the same six mice per timepoint, *i.e.* the muscles were matched for each animal. If the ANOVA indicated a significant difference, Bonferroni’s multiple comparisons test was performed to assess significance between pairwise combinations of all five muscles. Correlation was assessed using Pearson’s product moment correlation (**Figure 4** and **Supplementary Figure 3**, **4C**, **5**, **6B** and **6D**). When *P* < 0.05, a dashed line of best fit was plotted on the graph. A complete line was added when *P* < 0.00256, which is the Bonferroni correction-adjusted *P* value for an α of 0.05 when performing 20 associated tests (*i.e.* the number of tests used per timepoint). To calculate the percentage difference in morphological variables between timepoints (**Supplementary Table 4**), the mean value at P31-32 was divided by the mean at P7 and multiplied by 100. Two-way ANOVAs were performed for each morphological trait (test variables being age and muscle), followed by Sidak’s multiple comparisons test to assess whether there was a difference between timepoints in each muscle.

**Table 1.** Characteristics of the five wholemount muscles used in this study. See also **Figure 1**.

| Muscle | Body position | Innervating nerve | % fast twitch | Animal | Reference |
| --- | --- | --- | --- | --- | --- |
| Epitrochleoanconeus (ETA) | Forelimb | Radial | 90 | Mouse | (Bradley *et al.*, 1989) |
| Flexor digitorum brevis (FDB) | Hindlimb | Medial plantar | 95.6 | Mouse | (Tarpey *et al.*, 2018) |
| Lumbricals of the foot | Hindlimb | Medial/lateral plantar | 90 | Mouse | (I. C. Smith *et al.*, 2013) |
| Lumbricals of the hand | Forelimb | Median/ulnar | 100 | Mouse | (Russell *et al.*, 2015) |
| Transversus abdominis (TVA) | Thorax | Lower intercostal | 68 | Rat | (Delp and Duan, 1996) |

### 2.7 Principal component analysis

Principal component analysis (PCA) was used for dimensionality reduction of the datasets of P7 and P31-32 NMJ morphological variables by collapsing the variables into new ‘principal components’ created through linear combinations of the original datasets (**Supplementary Figure 1**). Data.table (https://CRAN.R-project.org/package=data.table) and tidyverse (Wickham *et al.*, 2019) packages were used to load the morphological data into R (version 3.6.2). PCA plots and loading analysis were created using the PCAtools package (https://github.com/kevinblighe/PCAtools) and ggplot2 (Wickham, 2016).

### 2.8 Clustering analysis

Morphological variables were converted to z-scores and samples clustered by Euclidean distance and hierarchal clustering. The complete linkage method was used to find breaks. Euclidean distance between two samples is the square root of the sum of squared differences between all z-score-scaled morphological variables. Complete clustering is a distance metric that defines the distance between clusters as the maximum possible distance between points in a cluster. Heat maps and clustering were created using the pheatmap package (https://CRAN.R-project.org/package=pheatmap).

## 3. Results and Discussion

### 3.1 Wholemount immunohistochemical analyses of mouse muscles and NMJs

Five mouse muscles from diverse body positions with varying innervating nerves and fast twitch fibre percentages (**Table 1**), all of which are also found in humans, were selected for morphological analysis of NMJs (**Figure 1**). The lumbricals of the hand mediate metacarpophalangeal joint flexion (*i.e.* paw clasping), are innervated by the median/ulnar nerve and are composed of ~100% fast twitch muscle fibres (**Figure 1A**) (Russell *et al.*, 2015). Consisting of ~90% fast twitch fibres and innervated by the radial nerve, the epitrochleoanconeus (ETA) is located on the medial surface of the upper forelimb and contributes to forearm supination (*i.e.* outward rotation of the palm) (**Figure 1B**) (Bradley *et al.*, 1989). Innervated by lower intercostal nerves, the transversus abdominis (TVA) is a postural muscle also involved in respiration that is found in the anterior abdominal wall consisting of ~68% fast twitch fibres (**Figure 1C**) (Delp and Duan, 1996; Murray *et al.*, 2014). Situated in the hindlimb feet, the flexor digitorum brevis (FDB) and lumbricals aid metatarsophalangeal joint extension (*i.e.* paw opening) and flexion (*i.e.* paw clasping), respectively (**Figure 1D**). The FDB consists of ~96% fast twitch fibres and is innervated by the medial plantar nerve (Tarpey *et al.*, 2018), whereas the feet lumbricals are ~90% fast twitch and innervated by both medial and lateral plantar nerves (I. C. Smith *et al.*, 2013). All five muscles possess the major advantage that they are thin and flat, and can, therefore, be dissected and immunohistochemically processed as wholemount preparations without the need for sectioning (**Figure 1A-D i-ii**). This permits visualisation of the entire innervation pattern of these muscles using combined neuron-specific antibodies against SV2/2H3 and AChR-targeting α-BTX (**Figure 1A-D iii**), and accurate evaluation of NMJ structure (**Figure 1A-D iv**) (Sleigh *et al.*, 2014a).

**Figure 1.**
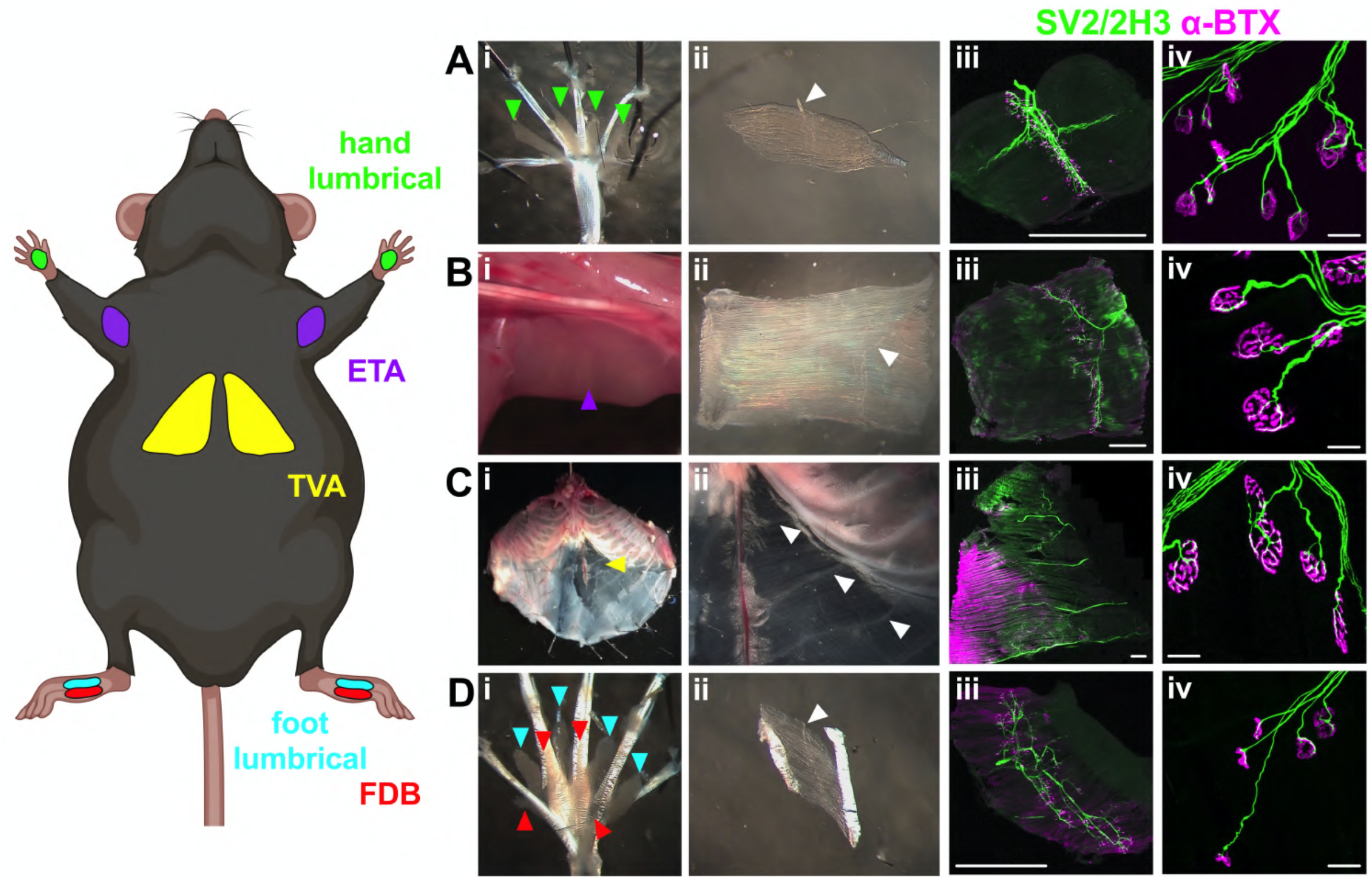
Dissection of five wholemount muscles for assessment of NMJ morphology. The lumbricals of the hand (green, **A**), the epitrochleoanconeus (ETA, purple, **B**), the transversus abdominis (TVA, yellow, **C**), the lumbricals of the foot (cyan, **D i**) and the flexor digitorum brevis (FDB, red, **D**) are five thin and flat muscles that can be dissected from mice (i and ii) and immunohistochemically analysed as wholemount preparations (iii) to assess pre-(SV2/2H3, green) and post-synaptic (alpha bungarotoxin, α-BTX, magenta) NMJ morphology (iv). Representative staining of feet lumbrical NMJs can be found in **Figure 2**. Muscles imaged for this figure were taken from a mouse at P31. Individual muscles in i panels are highlighted by arrowheads coloured the same as cartoon muscles in the mouse schematic on the left; this colour scheme is maintained throughout all figures and supplementary figures. White arrows in ii panels identify innervating nerves. Scale bars = 1 mm and 20 μm in iii and iv panels, respectively. The mouse schematic was created with BioRender (https://biorender.com). See also **Table 1**.

### 3.2 Mature post-synaptic NMJ morphology varies more than pre-synaptic between muscles

Muscles were dissected from wild-type mice aged P31-32, a timepoint chosen to represent a mature, early adult NMJ. To confirm this, NMJ development was assessed by calculating the percentage of synapses innervated by more than one motor neuron (*i.e.* polyinnervation), which is a key feature of immature NMJs (Sanes and Lichtman, 1999). As expected, all muscles had undergone the process of synapse elimination to become monoinnervated (**Figure 2A i**), while post-synapses displayed clear pretzel-like structures (**Figure 1A-D iv**). The foetal to adult AChR subunit switch is also usually complete by this stage (Missias *et al.*, 1996). Together, this suggests that analysis at P31-32 allows assessment of relatively well developed NMJs across all five muscles.

**Figure 2.**
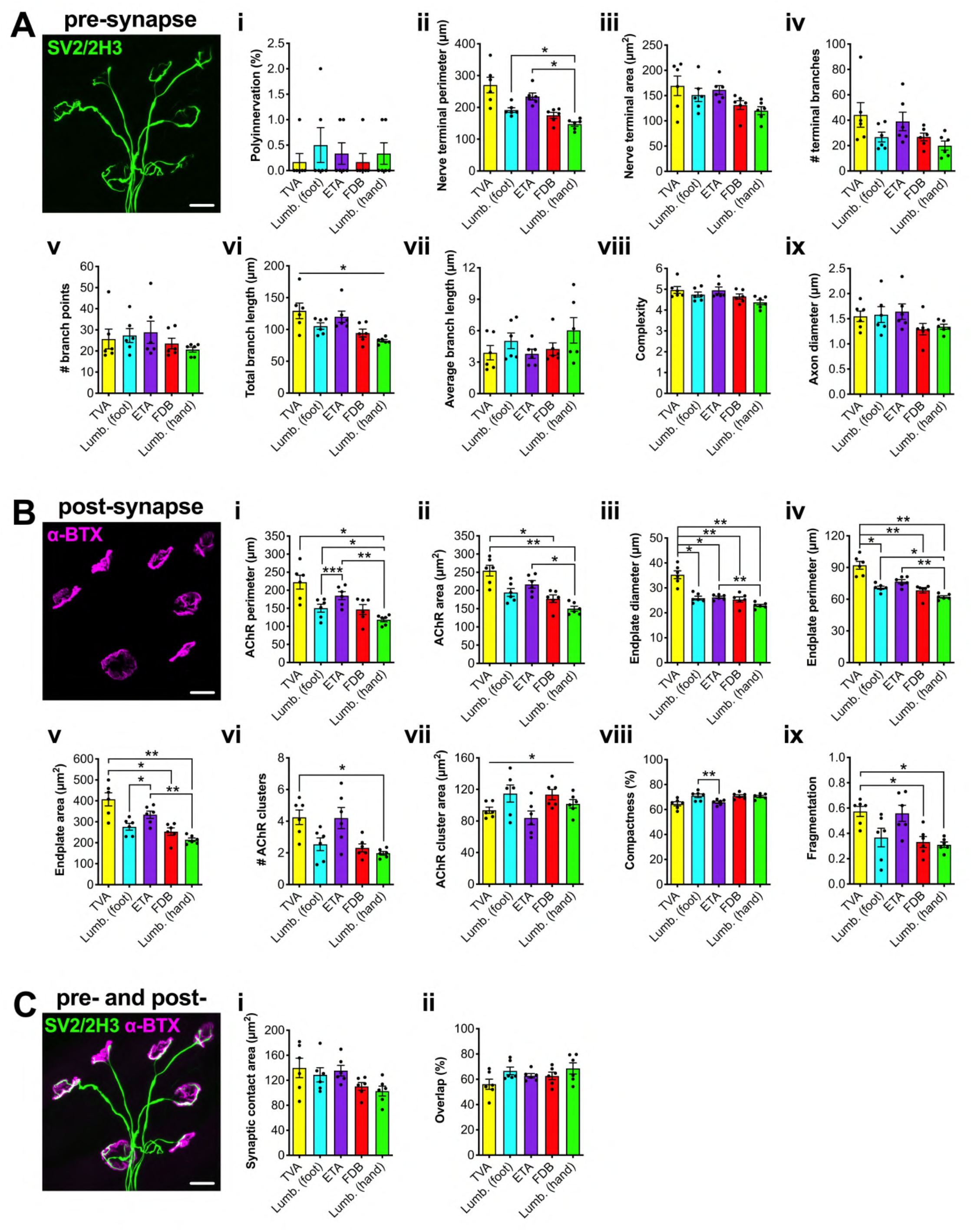
Post-synaptic NMJ morphology shows greater variability between muscles than pre-synaptic at maturity. Five different wholemount muscles were dissected from P31-32 mice and immunohistochemically processed to visualise pre- (**A**, SV2/2H3, green) and post-synaptic (**B**, α-BTX, magenta) NMJ morphology. The NMJ-morph workflow was followed to assess eight pre-synaptic variables (**A ii-ix**), nine post-synaptic variables (**B i-ix**), and two combined pre- and post-synaptic variables (**C i-ii**), details of which can be found in the Materials and Methods. Polyinnervation was scored separately by eye (**A i**). The representative immunofluorescence image was taken of a foot lumbrical muscle dissected at P31. Individual variables were analysed using a repeated-measures one-way ANOVA, details of which are provided in **Supplementary Table 2**. * *P* < 0.05, ** *P* < 0.01, *** *P* < 0.001 Bonferroni’s multiple comparisons test. Means ± standard error of the mean are plotted (*n* = 6), as well as individual data points generated from each individual mouse (*i.e.* the mean values from 20 NMJs). Lumb. (foot), lumbricals of the foot; Lumb. (hand), lumbricals of the hand. Scale bars = 20 μm. See also **Table 2** and **Supplementary Table 2**.

NMJ-morph was then used to assess pre-synaptic (**Figure 2A**), post-synaptic (**Figure 2B**) and overlapping (**Figure 2C**) features of NMJ morphology (**Table 2**, **Supplementary Table 2**). In all graphs, muscles are plotted on the x-axis in ascending order of the percentage of fast twitch fibres. Of eight pre-synaptic variables, only nerve terminal perimeter showed significant differences between individual muscles, with lumbricals of the hand having smaller perimeters than foot lumbricals and the ETA (**Figure 2A ii**). The TVA had the largest average nerve terminal perimeter and the hand lumbricals the smallest, although nerve terminal area, complexity, and axon diameter were all similar across muscles. Total branch length showed a significant difference using a repeated-measures one-way ANOVA, yet no inter-muscle differences were subsequently identified (**Figure 2A vi**). As a relative measure of the morphological range observed between muscles, the fold difference between the maximum and minimum values for each variable were calculated (**Table 2**). The number of terminal branches showed the greatest fold difference (2.21), while complexity displayed the least (1.14).

**Table 2.** P31-32 NMJ morphological variables. The mean values for all NMJ variables were calculated for each muscle per animal from 20 NMJs, the means of which (*n* = 6) were then recorded for each muscle. The plotted mean, maximum (Max) and minimum (Min) values for each NMJ variable were determined from these calculated values. To gauge the range in morphology observed between muscles, the fold difference (Fold Diff.) was calculated by dividing the maximum value by the minimum. Variables are divided into pre-synaptic at the top, post-synaptic in the middle, and combined pre- and post-synaptic at the bottom. See also **Figure 2** and **Supplementary Table 2**.

| Variable | Mean | Max | Min | Fold Diff. |
| --- | --- | --- | --- | --- |
| Polyinnervation (%) | 0.30 | 0.50 | 0.17 | 2.94 |
| Nerve terminal perimeter (μm) | 203.80 | 270.39 | 147.61 | 1.83 |
| Nerve terminal area (μm<sup>2</sup>) | 146.62 | 169.33 | 120.11 | 1.41 |
| # terminal branches | 31.32 | 44.09 | 19.98 | 2.21 |
| # branch points | 25.22 | 28.84 | 20.80 | 1.39 |
| Total branch length (μm) | 106.28 | 129.27 | 82.31 | 1.57 |
| Average branch length (μm) | 4.61 | 6.03 | 3.80 | 1.59 |
| Complexity | 4.74 | 4.97 | 4.37 | 1.14 |
| Axon diameter (μm) | 1.48 | 1.64 | 1.29 | 1.27 |
| AChR perimeter (μm) | 164.56 | 222.39 | 118.45 | 1.88 |
| AChR area (μm<sup>2</sup>) | 198.64 | 254.43 | 150.07 | 1.70 |
| Endplate diameter (μm) | 27.14 | 35.28 | 22.92 | 1.54 |
| Endplate perimeter (μm) | 74.04 | 92.07 | 62.22 | 1.48 |
| Endplate area (μm<sup>2</sup>) | 296.98 | 407.30 | 214.33 | 1.90 |
| # AChR clusters | 3.06 | 4.25 | 1.98 | 2.15 |
| AChR cluster area (μm<sup>2</sup>) | 101.52 | 114.73 | 83.94 | 1.37 |
| Compactness (%) | 68.57 | 71.02 | 64.60 | 1.10 |
| Fragmentation | 0.43 | 0.58 | 0.31 | 1.86 |
| Synaptic contact area (μm<sup>2</sup>) | 123.34 | 139.75 | 102.20 | 1.37 |
| Overlap (%) | 63.36 | 68.57 | 56.11 | 1.22 |

Contrasting with the nerve, the post-synapse displayed many more significant differences in morphology between muscles, with only AChR cluster area not doing so, albeit with significance indicated by an ANOVA (**Figure 2B vii**). The TVA had the largest reported values for most post-synaptic variables and hand lumbricals the smallest. The number of AChR clusters showed the greatest fold difference (2.15) and compactness the smallest (1.10) (**Table 2**). Finally, there were no significant differences in the two combined pre- and post-synaptic variables of synaptic contact area and overlap (**Figure 2C**), with maximum to minimum fold differences of 1.37 and 1.22, respectively (**Table 2**). This indicates that despite morphological variety between muscles, the level of nerve to muscle interaction remains fairly constant, suggesting that pathological denervation assessed in different muscles can be reliably compared.

As twenty separate repeated-measures ANOVAs were conducted, the Bonferroni correction was subsequently applied to determine that a *P* value of less than 0.00256 for each test is required to maintain an α of 0.05. With this more stringent *P* value, no significant differences were observed in pre-synaptic or combined morphological variables, whereas six post-synaptic variables remained significant (**Supplementary Table 2**). These data indicate that mature neuromuscular synapses found in a diverse set of wholemount muscles display significant variations in their structure that are observed consistently in their post-synapse.

### 3.3 Pre- and post-synaptic architecture are more varied at immature NMJs

To determine whether developing NMJs display similar distinctions in NMJ architecture, morphological analysis was also performed at P7. Polyinnervation counts confirmed that the process of synapse elimination was not yet complete in any of the five wholemount muscles (**Figure 3A i**), indicating that the P7 timepoint allows analysis of immature NMJs. It has been suggested previously that synapse elimination generally occurs along an anterior-to-posterior gradient, with NMJs further away from the thorax taking longer to undergo this important developmental process (Bixby and van Essen, 1979). However, the TVA, which is the muscle closest to the thorax of the five assessed here, showed the highest percentage of polyinnervated NMJs (~36%). Moreover, the three muscles in the paws showed very similar degrees of polyinnervation (18-19%) despite the FDB and foot lumbricals being more distal than hand lumbricals. These data indicate that in P7 mice there is no body-wide location effect on synapse elimination. Nonetheless, regional distinctions in polyinnervation are known (Bixby and van Essen, 1979), so perhaps additional factors, such as muscle fibre type (Lee, 2019), contribute to the observed results (see below).

**Figure 3.**
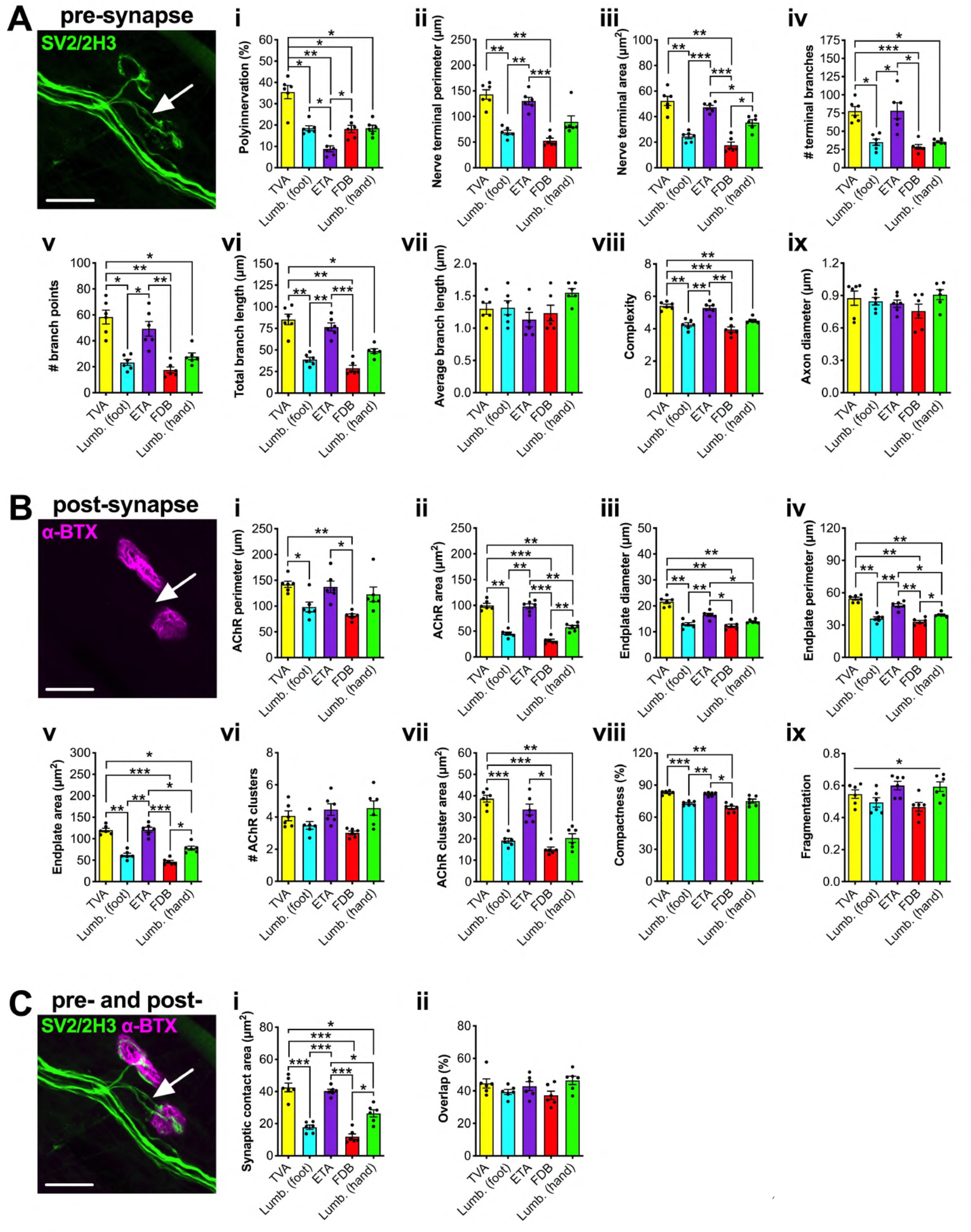
The developing NMJ shows greater morphological variability between muscles than mature synapses. Five different wholemount muscles were dissected from P7 mice and immunohistochemically processed to visualise pre- (**A**, SV2/2H3, green) and post-synaptic (**B**, α-BTX, magenta) NMJ morphology. The NMJ-morph workflow was followed to assess eight pre-synaptic variables (**A ii-ix**), nine post-synaptic variables (**B i-ix**), and two combined pre- and post-synaptic variables (**C i-ii**), details of which can be found in the Materials and Methods. Polyinnervation was scored separately by eye (**A i**). The representative immunofluorescence image was taken using a TVA muscle. The arrow identifies a polyinnervated NMJ. Individual variables were analysed using a repeated-measures one-way ANOVA, details of which are provided in **Supplementary Table 3**. * *P* < 0.05, ** *P* < 0.01, *** *P* < 0.001 Bonferroni’s multiple comparisons test. Means ± standard error of the mean are plotted (*n* = 6), as well as individual data points generated from each individual mouse (i.e. the mean values from 20 NMJs). Lumb. (foot), lumbricals of the foot; Lumb. (hand), lumbricals of the hand. Scale bars = 20 μm. See also **Table 3** and **Supplementary Table 3**.

The NMJ-morph software was used to assess pre-synaptic (**Figure 3A**), post-synaptic (**Figure 3B**) and overlapping (**Figure 3C**) morphologies at the developing NMJ (**Table 3**, **Supplementary Table 3**). Unlike at mature synapses, all but two variables, average branch length (**Figure 3A vii**) and axon diameter (**Figure 3A ix**), showed significant distinctions between muscles at P7, which all remained significant with the Bonferroni correction applied (**Supplementary Table 3**). Reflective of several pre-synaptic features at P31-32, the TVA and ETA values were larger than FDB and foot lumbricals for all other pre-synaptic variables, with the hand lumbricals showing intermediate values. The number of branch points displayed the greatest fold difference between maximum and minimum values (3.30), while axon diameter displayed the least (1.20) (**Table 3**). The developing post-synapse showed a similar number of significant differences between muscles as the mature NMJ, with all variables showing distinctions, except for the number of AChR clusters (**Figure 3B vi**).

**Table 3.** P7 NMJ morphological variables. The mean values for all NMJ variables were calculated for each muscle per animal from 20 NMJs, the means of which (*n* = 6) were then recorded for each muscle. The plotted mean, maximum (Max) and minimum (Min) values for each NMJ variable were determined from these calculated values. To gauge the range in morphology observed between muscles, the fold difference (Fold Diff.) was calculated by dividing the maximum value by the minimum. Variables are divided into pre-synaptic at the top, post-synaptic in the middle, and combined pre- and post-synaptic at the bottom. See also **Figure 3** and **Supplementary Table 3**.

| Variable | Mean | Max | Min | Fold Diff |
| --- | --- | --- | --- | --- |
| Polyinnervation (%) | 19.90 | 35.50 | 8.83 | 4.02 |
| Nerve terminal perimeter (μm) | 96.92 | 142.96 | 52.91 | 2.70 |
| Nerve terminal area (μm<sup>2</sup>) | 35.39 | 52.27 | 17.63 | 2.96 |
| # terminal branches | 51.05 | 78.25 | 28.57 | 2.74 |
| # branch points | 35.38 | 58.31 | 17.68 | 3.30 |
| Total branch length (μm) | 55.66 | 85.26 | 28.80 | 2.96 |
| Average branch length (μm) | 1.31 | 1.55 | 1.15 | 1.35 |
| Complexity | 4.68 | 5.42 | 3.96 | 1.37 |
| Axon diameter (μm) | 0.84 | 0.91 | 0.76 | 1.20 |
| AChR perimeter (μm) | 116.44 | 142.25 | 81.63 | 1.74 |
| AChR area (μm<sup>2</sup>) | 66.62 | 100.06 | 31.66 | 3.16 |
| Endplate diameter (μm) | 15.48 | 21.63 | 12.33 | 1.75 |
| Endplate perimeter (μm) | 42.31 | 54.55 | 33.06 | 1.65 |
| Endplate area (μm<sup>2</sup>) | 85.59 | 121.07 | 46.33 | 2.61 |
| # AChR clusters | 3.92 | 4.56 | 3.02 | 1.51 |
| AChR cluster area (μm<sup>2</sup>) | 25.38 | 38.70 | 15.10 | 2.56 |
| Compactness (%) | 76.22 | 82.94 | 68.72 | 1.21 |
| Fragmentation | 0.54 | 0.60 | 0.47 | 1.28 |
| Synaptic contact area (μm<sup>2</sup>) | 27.77 | 42.53 | 11.88 | 3.58 |
| Overlap (%) | 42.11 | 46.45 | 37.31 | 1.24 |

When the Bonferroni correction was applied, AChR area and endplate diameter, perimeter and area repeated as being significantly different, whereas AChR perimeter and fragmentation did not (**Supplementary Table 3**). However, AChR cluster area and compactness were significantly different at this timepoint. Similar to pre-synaptic morphology, the TVA and ETA generally had the largest post-synaptic structures, followed by the hand lumbricals then the FDB and foot lumbricals. AChR area showed the greatest fold difference between muscles (3.16), while compactness showed the smallest (1.21) (**Table 3**). Finally, consistent with the myriad of pre- and post-synaptic distinctions between muscles, the synaptic contact area showed significant variation (**Figure 3C i**, **Supplementary Table 3**) and a fold difference between maximum and minimum values of 3.58 (**Table 3**). Repeating the finding at P31-32, the percentage overlap between pre- and post-synapse did not differ (1.24 fold difference) (**Figure 3C ii**), once again confirming the possibility of cross-muscle denervation analysis. Together, these analyses of the developing NMJ indicate that there is greater variation between muscles at P7 than at the mature adult synapse; the post-synapse displays a comparable level of inter-muscle variation at both timepoints, while the pre-synapse shows greater irregularity during immaturity.

### 3.4 NMJ endplate area is a powerful discriminator of developmental stage and muscle type

To understand which morphological variables changed most throughout NMJ development, a principal component analysis (PCA) was performed. PCA is an exploratory data analysis tool widely used for dimensionality reduction because of its ability to condense large datasets in an interpretable way. The technique functions by collapsing the existing variables into new principal components (PCs) that are created through linear combinations of the original dataset. During the creation of PCs, the variance between components is maximised so that each new component is as different from each other as possible, and thus generates additional information about the original dataset.

NMJs from both P7 and P31-32 were plotted along the first, and most important, two PCs (**Supplementary Figure 1A**). PC1 accounts for 90.52% of the variation in the dataset and clearly separated the two timepoints, with some muscles within timepoints also diverging (e.g. P7 FDB and TVA). PC2 explains only 4.87% of the variation within the dataset, but roughly separated NMJs across different muscles, with little to no separation of timepoints. Distribution of muscles across PC1 and PC2 indicates that at P31-32, the hand lumbricals are most different from the TVA, while FDB and foot lumbricals are most similar (**Supplementary Figure 1A**), which are corroborated by clustering analysis (**Supplementary Figure 1B**) and pairwise significance testing between all muscles for all variables (**Supplementary Table 2**). At P7, the FDB and foot lumbricals are again most similar, while the TVA and ETA are most dissimilar to the FDB and foot lumbricals (**Supplementary Figure 1A**), which are again supported by clustering analysis (**Supplementary Figure 1C**) and pairwise significance testing (**Supplementary Table 3**). Generally speaking, NMJs at maturity were more variable than developing NMJs (greater spread of datapoints), especially along PC2, with the TVA showing the greatest irregularity at P31-32 and the ETA at P7. These patterns were corroborated by the clustering analyses (**Supplementary Figure 1B, C**). This variability may be linked with fibre type percentages and sampling of imaged NMJs, as the TVA, which has the greatest variability at maturity, has the highest percentage of slow twitch fibres (~32%). This pattern is not observed at P7, but muscle fibre types are not yet fully determined at this age in mice (Schiaffino and Reggiani, 2011). That being said, the 100% fast twitch hand lumbrical muscle shows similar variability at P31-32, if not more, to the remaining three muscles. The variability may thus instead be linked to additional properties of the muscle, e.g. variability in fibre or overall muscle sizes.

To find which features contributed most to PC1 and PC2, a loading plot analysis was performed (**Supplementary Figure 1D**), which shows how strongly each morphological variable contributes to each PC. Endplate area contributed the most to PC1, with nerve terminal area, AChR perimeter and AChR cluster area also adding. The endplate area strongly reflects the overall size of the NMJ because it includes the area contained within the perimeter of α-BTX staining. Analysis of NMJs in nine different muscles from wild-type CD1 mice aged six weeks corroborate this finding that NMJ size drives the majority of variation between different muscles (Jones *et al.*, 2016). The number of terminal branches and AChR perimeter contributed the most to PC2, with much smaller influences from polyinnervation and compactness.

Finally, an Eigencor plot was created, which depicts Pearson product moment correlation coefficients (*r*) between PCs and the four metadata variables of percentage fast twitch muscle fibres, animal weight, sex, and age (**Supplementary Figure 1E**). PC1 correlated significantly with age (*P* < 0.001, *r* = 0.87), which is unsurprising as this PC clearly separated developmental and mature NMJs, and weight (*P* < 0.001, *r* = −0.84), which is obviously impacted by age (**Supplementary Table 1**). Nevertheless, samples within timepoints did not appear to cluster by weight (**Supplementary Figure 1B, C**), indicating that small differences in body weight at a particular timepoint do not greatly impact NMJ morphology. Age and weight also correlated with PC2, although less so (*P* < 0.05). However, muscle fibre type correlated most significantly with PC2 (*P* < 0.001, *r* = −0.60), which indicates that fibre type percentage is possibly driving distinctions in NMJ morphology between the assessed muscles. No correlations were observed between sex and PCs suggesting that up to the age of P31-32, there are no major differences between NMJs of male and female mice, which was corroborated by the clustering analyses (**Supplementary Figure 1B, C**). However, as the two sexes diverge in muscle mass and strength later in life, this may not continue to be observed.

### 3.5 Morphological change over NMJ development differs between muscles

As there is more structural variety at developing NMJs than mature synapses, the rate of maturation across the muscles is likely to differ, such that differences at P7 become smaller by P31-32. Supporting this is the observation that synapse elimination does not occur to the same extent by P7 in all five muscles (**Figure 3A i**). To assess morphological development and growth at the NMJ, variables from both timepoints were re-plotted on the same graphs (**Supplemental Figure 2**) and the percentage change over time calculated for each muscle (**Supplementary Table 4**). Two-way ANOVAs were performed on all 20 NMJ morphological properties, with age and muscle being the two statistical test variables. Applying the Bonferroni correction for using 20 associated tests, all morphological variables except for complexity showed a significant difference between ages (**Supplementary Table 4**). There is also a significant difference between muscles for most variables (with Bonferroni correction applied), except for average branch length, axon diameter, AChR cluster area, compactness, and overlap (**Supplementary Table 4**). Similarly, individual muscles show differences in most variables between timepoints, with at least one muscle displaying a difference in every variable (**Supplementary Figure 2**). Additionally, several different pre- and post-synaptic variables show a significant interaction, indicating that the amount of change occurring at the NMJ during post-natal development is muscle-specific, *i.e.* NMJs across different muscles do not develop and grow at the same rate between P7 and P31-32. Indeed, lumbricals of the foot and FDB showed the greatest average change per morphological variable (260% and 303%, respectively), while the ETA (168%) and TVA (170%) the smallest (values include polyinnervation; **Supplementary Table 4**).

These differences in development/growth rate perhaps reflect the function of each muscle and their relative importance during early post-natal life. The TVA and ETA are the two muscles with the largest NMJs at both timepoints and that display the least change from P7 to P31-32. These two muscles are therefore perhaps some of the most well developed during the first week of life in mice, at least in terms of the size of pre- and post-synaptic structures (keeping in mind that polyinnervation is rife in the TVA at P7). The TVA has a role in respiration (Mesquita Montes *et al.*, 2016), so has a clear need to be functional from birth. Furthermore, feeding behaviours of pups are likely to be more dependent on postural muscles and those that mediate larger movements of the limbs than perhaps those that facilitate finer movements of the paws. Being able to breathe and feed immediately after birth are clearly two of the most important early mouse behaviours, so perhaps that is why the TVA and ETA muscles appear most well developed and why muscles generally show distinctions in their rates of NMJ development and growth.

### 3.6 Fast twitch fibres are more likely to be innervated by smaller NMJs

A wealth of data on NMJ morphology has now been generated at two different timepoints in five different mouse muscles, with a plethora of distinctions observed between muscles (**Figure 2** and **3**) and timepoints (**Supplementary Figure 2**). A possible cause for these differences is the relative abundance of fast and slow twitch fibres. Indeed, NMJ size has been previously reported to be related to muscle fibre type (Prakash *et al.*, 1996; Wood and Slater, 1997), and PC2, which separated NMJs across muscle types, correlated significantly with the percentage of fast twitch muscle fibres (**Supplementary Figure 1E**). Thus, all NMJ variables at P7 and P31-32 were correlated with previously reported (**Table 1**) fast twitch muscle fibre percentages (**Figure 4**, **Supplementary Table 5**).

**Figure 4.**
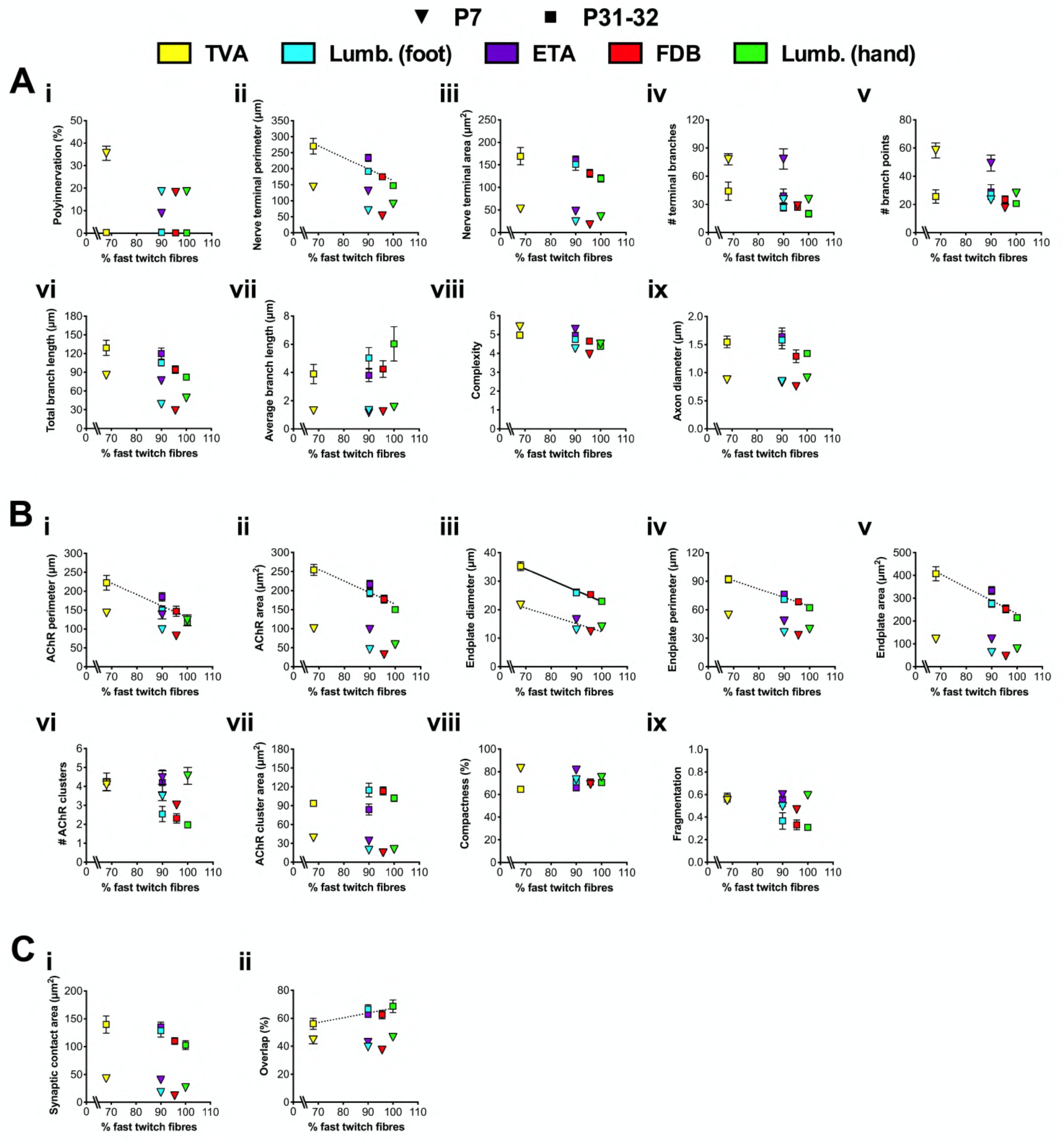
Mature post-synaptic morphology correlates with muscle fibre type indicating that smaller NMJs innervate fast twitch fibres. The percentage of fast twitch muscle fibres found within the five dissected wholemount muscles (see **Table 1**) was correlated with all twenty pre-synaptic (**A**), post-synaptic (**B**), and combined (**C**) NMJ morphological variables from both P7 (inverted triangles) and P31-32 (squares) timepoints. Correlation was assessed by calculating Pearson’s product moment correlation coefficient (*r*), the results of which are presented in **Supplementary Table 5** along with associated *P* values. When the *P* value was < 0.05, a dashed line of best fit was plotted on the graph. A complete line was added when the *P* value was < 0.00256, which is the Bonferroni correction-adjusted *P* value for an α of 0.05 when performing 20 associated tests. Means ± standard error of the mean are plotted for each NMJ variable (*n* = 6). Lumb. (foot), lumbricals of the foot; Lumb. (hand), lumbricals of the hand. See also **Supplementary Table 5**.

First, assessing synapse elimination, no correlations were observed between the percentage of fast twitch fibres and polyinnervation at both timepoints (**Figure 4A i**). This contrasts with the findings of Lee (2019), who showed a significant correlation between slow twitch fibres and polyinnervation across EDL, soleus, and sternomastoid muscles of P9 wild-type mice. This was directly corroborated by observations of individual fibres within the soleus muscle indicating a greater degree of polyinnervation at NMJs on slow fibres (Lee, 2019). The conflict between the results obtained in this and the previous study could be due to several reasons, including the use of different timepoints and muscles. Several additional publications found no link between muscle fibre type and synapse elimination, albeit in different species (mouse vs. rabbit) and using different methods (direct vs. indirect) (Bixby and van Essen, 1979; Cramer and Van Essen, 1995; Soha *et al.*, 1987). Nevertheless, assessing neuronal inputs at the NMJ in combination with immunofluorescent fibre typing is a powerful tool to better understand the importance of fibre type on synapse elimination and should therefore be used in the future to address this within the five wholemount muscles.

Moving on to the influence of fibre type on additional pre-synaptic (**Figure 4A**), post-synaptic (**Figure 4B**) and overlapping (**Figure 4C**) variables (**Supplementary Table 5**), no significant correlations were observed between the percentage of fast twitch muscle fibres and any of the pre-synaptic morphologies. Without Bonferroni correction, nerve terminal perimeter at P31-32 was significantly inversely correlated (*P* = 0.031), such that higher fast twitch fibre percentages are linked to smaller perimeters (**Figure 4A ii**); however, the *P* value did not reach significance when the twenty statistical tests were taken into account. In contrast, five out of nine of the post-synaptic variables were inversely correlated without use of Bonferroni correction (**Figure 4B i-v**), of which endplate diameter remained significantly associated with fibre type when using the more stringent *P* value cut-off (**Figure 4B iii**). The two variables showed an inverse correlation, indicating that smaller endplate diameters are linked with fast twitch muscle fibres. Of all the variables at P7, only endplate diameter again inversely correlated, albeit not when the Bonferroni correction was applied. Finally, of the two combined variables, overlap percentage significantly correlated with fibre type (**Figure 4C ii**), although, again, this was not significant when the Bonferroni correction was applied. Correlation between fibre type percentages and the rate of NMJ development/growth between P7 and P31-32 in all twenty morphological variables was also assessed, but no significant associations were observed (**Supplementary Figure 3**, **Supplementary Table 6**). This suggests that fibre type has little to no influence on change in NMJ morphology over the first month of post-natal development. Instead, perhaps one or a combination of the myriad transmembrane receptors, secreted ligands and other trans-synaptic proteins found at the NMJ are driving these inter-muscle distinctions, several of which are known to be involved in the developmental process of synapse elimination (English, 2003; Singhal and Martin, 2011).

Consistent with PC2 of the PCA, muscle fibre type does indeed appear to influence aspects of NMJ morphology, with smaller endplate diameters being significantly associated with fast twitch fibres (**Figure 4B iii**). Although all other parameters at both time points did not significantly correlate with fibre type, many showed the same general trend as endplate diameters, *i.e.* larger features of NMJ morphology tended to be observed in muscles with lower percentages of fast twitch fibres, and this was particularly true for the post-synaptic architecture. There are studies in individual muscles that both support (McArdle and Sansone, 1977; Wood and Slater, 1997) and contrast with (Prakash *et al.*, 1996) this finding; however, NMJ size has been reported to correlate with muscle fibre size in several different species (Balice-Gordon and Lichtman, 1990; Harris, 1954; Kuno *et al.*, 1971; Nyström, 1968), which may be contributing to the presented findings.

The five wholemount muscles were therefore re-mounted after teasing so that muscle fibre diameters could be assessed (**Supplementary Figure 4A**). Only muscles from the P31-32 timepoint were used because correlations with fibre type were more prevalent than at P7 (**Figure 4**). Significant differences in fibre diameter between several muscles were observed, with the ETA having the largest fibres and the hand lumbricals the smallest (2.05 fold difference) (**Supplementary Figure 4B**). When fibre diameters were correlated with the percentage of fast twitch fibres, no significant relationship was found (**Supplementary Figure 4C**). This indicates that fibre type does not impact fibre diameter across the five wholemount muscles that were analysed. This contrasts with the general assumption that fast twitch fibres have larger diameters (Stifani, 2014), and indicate that subsets of muscles may display idiosyncratic fibre features, perhaps linked with overall size and thickness of the muscle, as has been observed previously (Alnaqeeb and Goldspink, 1987). To determine whether fibre diameters may be confounding the observed link between fibre type and post-synaptic architecture (**Figure 4**), fibre diameters were correlated with all NMJ morphological variables at P31-32 (**Supplementary Figure 5**, **Supplementary Table 7**). No significant associations were observed, suggesting that fibre diameter does not have a major influence on NMJ morphology, with the major caveat that the individual fibres that were measured were not the same ones from which NMJs were studied. Similar to the polyinnervation phenotype, future work is therefore needed to immunohistochemically co-assess fibre types, their diameters and associated NMJ morphologies within the five muscles to address this more directly. This will be difficult to perform for all fibre subtypes, as the MHC-specific antibodies work only on non-fixed muscles (Rossor *et al.*, 2020), while fixed samples are needed for SV2/2H3 staining; that being said, co-staining of fixed muscle with a slow fibre marker is possible (Lee, 2019).

When normalised to muscle fibre size, it has been shown that slow twitch NMJs in rat diaphragm muscle cover a relatively greater area than fast twitch NMJs (Prakash *et al.*, 1996, 1995b; Prakash and Sieck, 1998). To determine whether a similar pattern is observed in mice, the ratios of P31-32 endplate diameters and endplate areas to fibre diameters were calculated for each muscle and correlated to fibre type (**Supplementary Figure 6**). These two morphological variables were chosen because endplate diameter significantly correlated with fibre type (**Figure 4B iii**), while endplate area drove most NMJ variation in PC1 (**Supplementary Figure 1D**) and is the best proxy for overall NMJ size. While significant differences were observed between muscles (**Supplementary Figure 6A** and **6C**), no correlation between fibre type and relative endplate diameter nor relative endplate area were observed (**Supplementary Figure 6B** and **6D**). This suggests that the pattern of NMJ size to fibre type ratio observed in rat diaphragm is not replicated in these mouse muscles, and that instead it is the post-synaptic morphology of mouse NMJs that is associated with muscle fibre type.

### 3.7 Limitations

While this study presents an unbiased assessment of neuromuscular synapse morphology in wild-type mice at early post-natal and adult stages, there are several limitations to this work that should be considered when interpreting the findings. The fibre type percentages were drawn from several different published studies (**Table 1**), which may subtly impact the consistency and accuracy of the fibre typing across all five muscles. Similarly, the percentage of fast twitch fibres in mouse TVA muscle could not be identified from the literature, hence the closest species available, the rat, was instead used. Not all muscles show similar slow and fast twitch fibre type percentages between these two rodents, e.g. the plantaris is 0% slow twitch in mouse and ~8% in rat, with rat muscles reported to broadly show a relatively “slower” phenotype than mouse muscles (Bloemberg and Quadrilatero, 2012), which is something generally observed in larger mammalian species (Schiaffino and Reggiani, 2011). The percentage of fast twitch fibres in the TVA reported in **Table 1** and upon which correlation analyses were performed may thus be an underestimate for mice. Nevertheless, several muscles between the two species are comparable, e.g. tibialis anterior, EDL, and masseter (Bloemberg and Quadrilatero, 2012; Gorza, 1990; Schiaffino, 1974). Furthermore, given that the TVA is at one end of the fibre type spectrum of the muscles analysed in this study, the impact on correlations of the mouse TVA having a different fast twitch percentage from rat should be less than if the TVA was found within the spectrum. In relation to this and the applicability of the findings of this study for human muscles, fibre type percentages are also only infrequently conserved between mice and humans (Schiaffino and Reggiani, 2011). An additional potential limitation is that, in the interest of time, 20 NMJs per muscle per timepoint were analysed rather than the suggested 30 (Jones *et al.*, 2016). Nonetheless, graphical evidence is provided that there is little to no distinction in NMJ variables when 20 to 50 NMJs were observed (Jones *et al.*, 2016). Consistent with this, limited variation was seen between mean morphological values within the five muscles across the six dissected mice at either timepoint. This is also pertinent to the point that several different hand lumbrical, feet lumbrical, and FDB muscles were combined in the presented analysis, because subtle distinctions in morphologies have been reported between the first and fourth lumbricals of the foot and the second and fourth lumbricals of the hands (Jones *et al.*, 2016). Smaller muscles were combined here because comparative denervation analyses have already been performed in these muscles in mouse models of neuromuscular disease (Sleigh *et al.* 2014b and unpublished work). To verify the presented findings, additional wholemount and thicker muscles of varying sizes should be analysed, including smaller predominantly slow twitch muscles and larger mainly fast twitch. Finally, it should be highlighted that in addition to fibre type differences between mice and humans, NMJs are also morphologically divergent, with human synapses being smaller, less complex, more fragmented and nummular (*i.e.* formed of coin-shaped patches), rather than pretzel-like, in shape (Jones *et al.*, 2017). Therefore, broad extrapolations from the mouse data generated here may be relevant to the human synapse, but finer, muscle-specific insights will be more tenuous.

## 4. Conclusion

Using an ImageJ-based workflow, morphometric data on mouse NMJs from five different wholemount muscles were generated at early post-natal and adult timepoints. Developing NMJs were shown to have much greater variability between different muscles in their pre- and post-synaptic structures than mature NMJs, which only showed distinctions in their post-synapse. It is well known that NMJ size and morphology are predictive of function and that muscle fibre types differ in their functional requirements of neurotransmission (Harris and Ribchester, 1979; Jones *et al.*, 2016; Kuno *et al.*, 1971; Ribchester *et al.*, 2004; Waerhaug and Lømo, 1994). Hence, NMJ morphologies at both timepoints were correlated with fast twitch muscle fibre percentages. Pre-synaptic neuronal structures did not associate with fibre type; however, there was a significant link between post-synaptic endplate diameter and fast twitch fibre percentage that was independent of muscle fibre diameter. This finding indicates that, in the murine muscles assessed, fast twitch fibres possess smaller NMJs, which may facilitate efficiency and speed of neurotransmission. Additionally, this study provides useful, non-biased, baseline data on the pre- and post-synaptic morphology of mouse NMJs in five anatomically and functionally diverse muscles at two different timepoints. These data can be used by the scientific community in future cross-study, comparative work assessing pathological changes occurring at the NMJ in mouse models of neuromuscular disease.

## Acknowledgements

The authors would like to thank Dr. David Villarroel-Campos (University College London) for critical comments on the manuscript, and the Schiavo and Linda Greensmith laboratories (University College London) for providing such a supportive, stimulating and collaborative environment in which to work. This work was supported by the Medical Research Council Career Development Award (MR/S006990/1) [JNS], a Wellcome Trust Senior Investigator Award (107116/Z/15/Z) [GS], the European Union’s Horizon 2020 Research and Innovation programme under grant agreement 739572 [GS] and a UK Dementia Research Institute Foundation award [GS]. The graphical abstract was created with BioRender.

## Author Contributions

JNS conceived the experiments; AM, JNS performed the research; AM, ALB, JNS analysed the data; GS provided expertise and discussion; AM, JNS wrote the manuscript with input from all authors. All authors approved submission of this work.

## Supplementary Figures

**Supplementary Figure 1.**
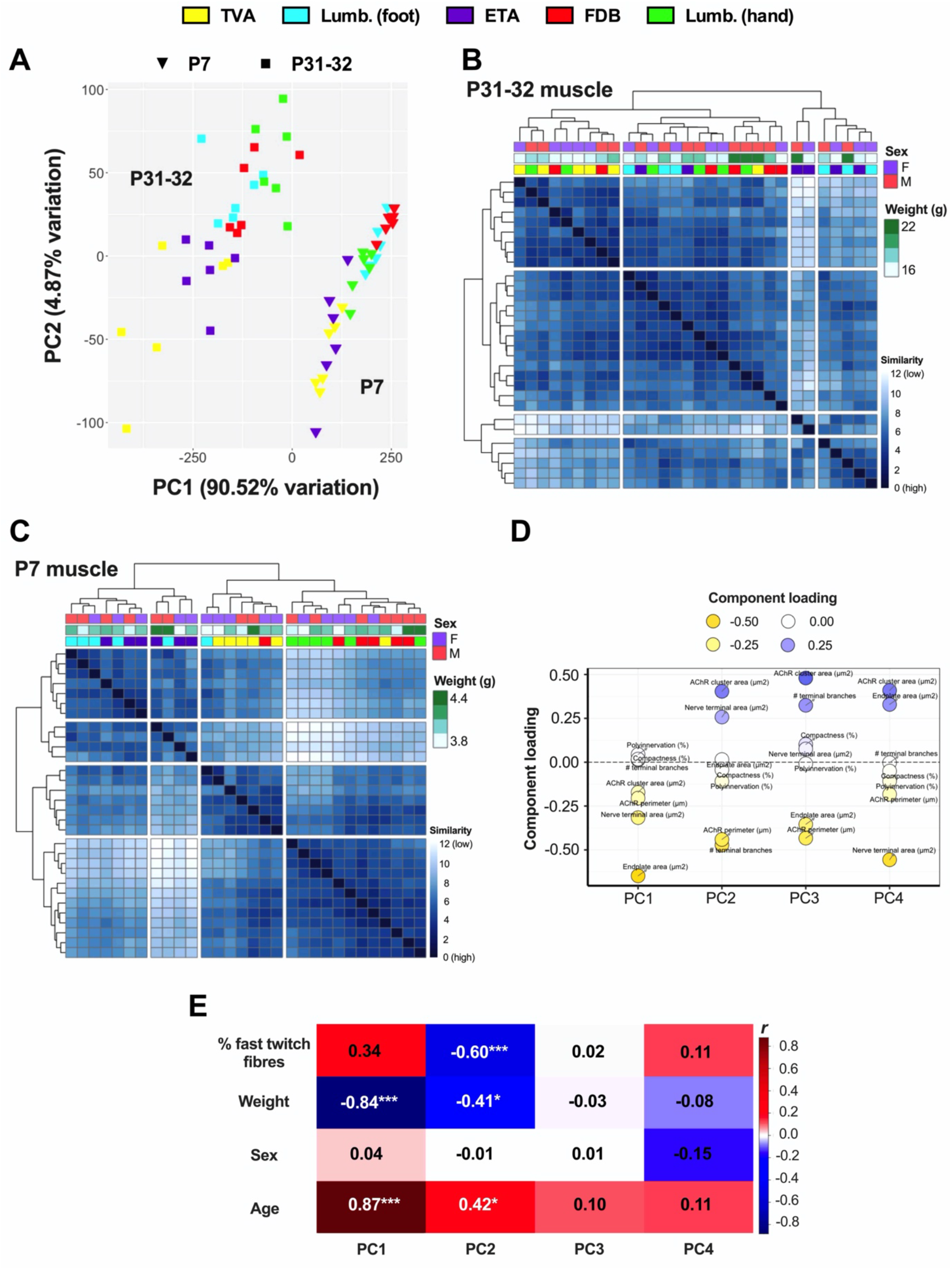
Principal components analysis and clustering analysis of NMJ morphology. (**A**) Two-dimensional PCA representation of NMJs as defined by 20 morphological variables (**Figure 2** and **3**) from each animal, colour-coded by muscle type (legend at top of the figure, and also used in B and C). Squares identify the P31-32 timepoint and inverted triangles the P7 timepoint. The x-axis represents the first principal component (PC1), accounting for 90.52% of the variability, and the y-axis, the second principal component (PC2), accounting for 4.87% of the variability. Each datum represents the centre point of the map positions of averages taken from 20 NMJs for each muscle from each animal. Note the ability of PC1 to distinguish between timepoints and PC2 to generally separate muscle types. Lumb. (foot), lumbricals of the foot; Lumb. (hand), lumbricals of the hand. (**B**, **C**) Heat maps generated by clustering analysis showing the sample similarly at P31-32 (B) and P7 (C) based on Euclidian distance. The dendrogram and breaks in the heat map represent hierarchal clustering of samples into k = 5 clusters by complete clustering. Each row and column represent a single muscle sample with the intensity of (blue) colour representing similarity between samples. Vertical length of the branches in the dendrogram give the distance between clusters. At P31-32, the TVA muscle is the most scattered throughout the cluster, with two of the TVA samples even forming a distinct cluster by themselves, suggesting that they are the most variable at this time point (B). At P7, most of the TVA and ETA muscles split off in the first branch in the clustering, suggesting these two muscles are most dissimilar from the other three. Note the lack of sample clustering based on sex and body weight at both timepoints. (**D**) PCA loadings plot identifying the morphological features that contribute most to PC1-PC4. Note that endplate area, nerve terminal area and AChR perimeter drive separation in PC1, whereas the number of terminal branches and AChR perimeter contribute most to PC2. PC3 and PC4 contribute little to the variability between samples (data not shown). (**E**) Eigencor plot identifying significant correlations between PC1-PC4 and metadata variables of ‘% fast twitch fibres’, ‘weight’, ‘sex’, and ‘age’. Pearson’s product moment correlation is reported on the plot, with r > 0 shaded red and r < 0 shaded blue. **\*** *P* < 0.05, **\*\*\*** *P* < 0.001 Bonferroni correction-adjusted *P* values.

**Supplementary Figure 2.**
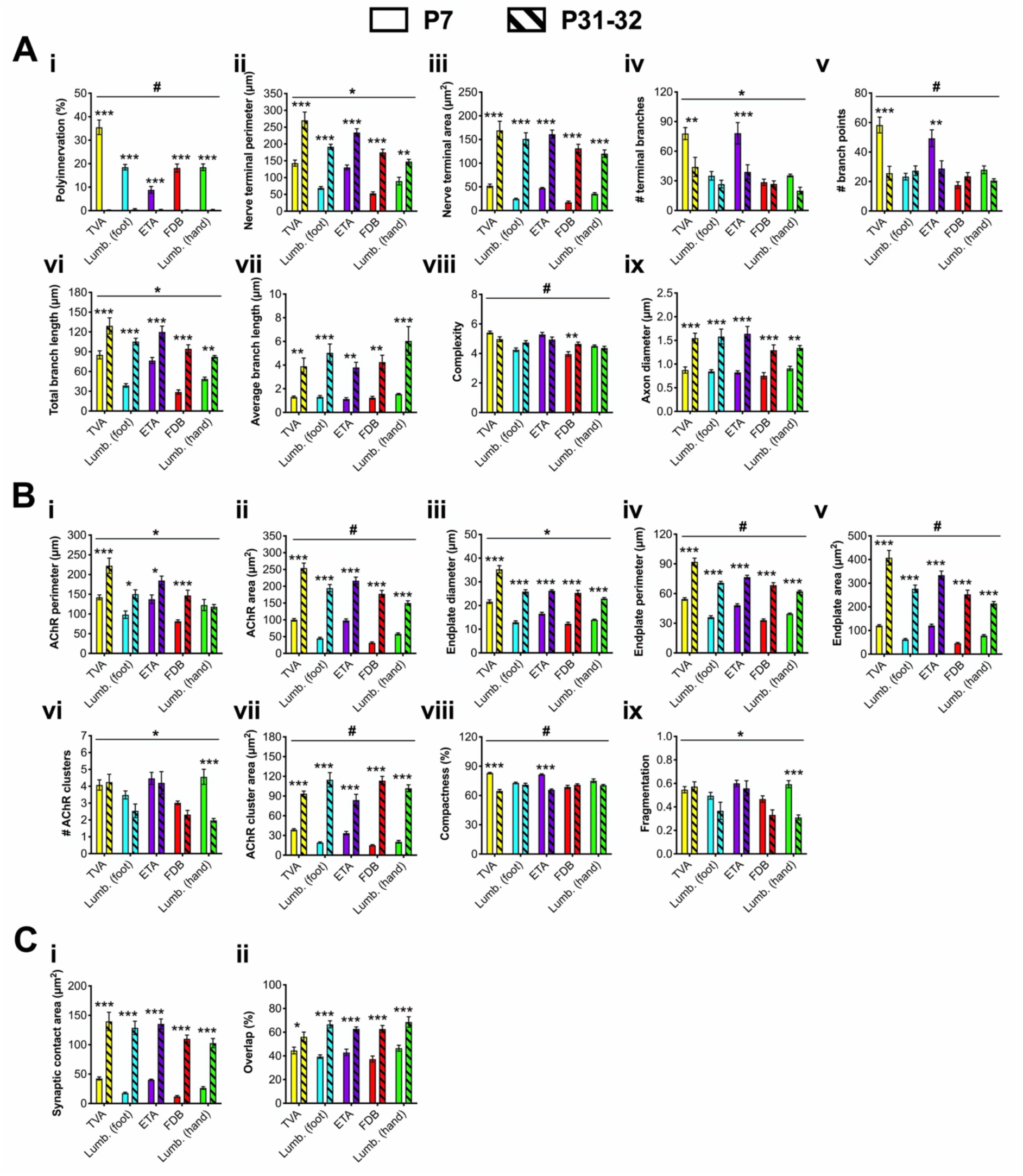
Morphological development of the NMJ varies between muscles. To assess the change in pre-synaptic (**A**), post-synaptic (**B**), and combined (**C**) morphological variables from P7 (clear bars) to P31-32 (hatched bars), data from each timepoint presented in **Figure 2** and **3** have been re-plotted on the same axes. Morphological variables were analysed using a two-way ANOVA (age and muscle as the test variables), details of which are provided in **Supplementary Table 4**, as are the percentage differences in each variable from P7 to P31-32. Interaction *P* values that are < 0.05 are denoted with a line and an asterisk at the top of the graph and those that are below < 0.00256 (*i.e.* the Bonferroni-corrected value for 20 associated tests) are identified with a ^#^. Means ± standard error of the mean are plotted (*n* = 6). **\****P* < 0.05, **\*\****P* < 0.01, **\*\*\****P* < 0.001 Sidak’s multiple comparisons test. Lumb. (foot), lumbricals of the foot; Lumb. (hand), lumbricals of the hand. See also **Supplementary Table 4**.

**Supplementary Figure 3.**
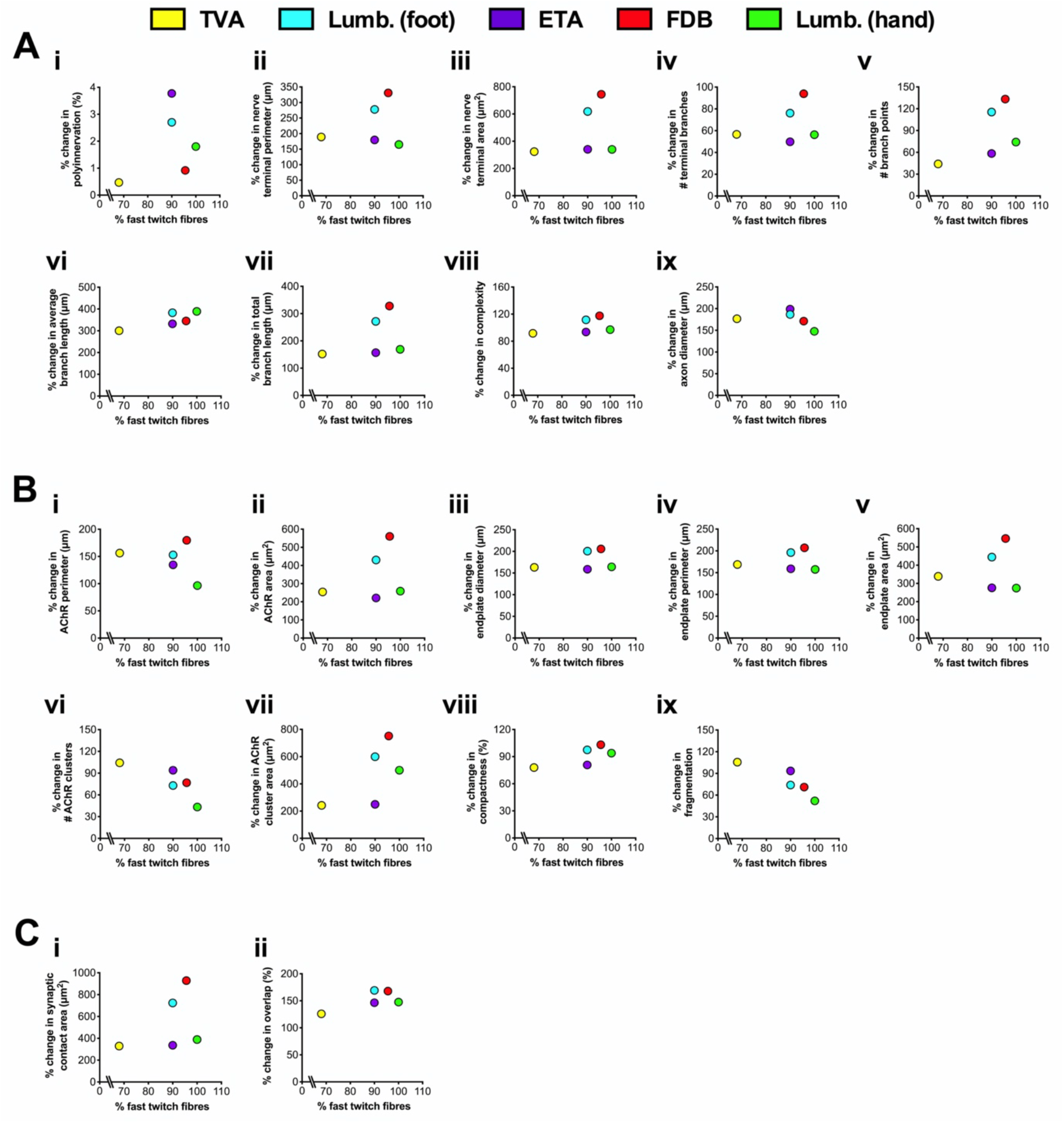
Muscle fibre type does not correlate with NMJ growth between post-natal developmental and early adulthood. The percentage of fast twitch muscle fibres found within the five dissected wholemount muscles (see **Table 1**) was correlated with the percentage change from P31-32 to P7 in all twenty pre-synaptic (**A**), post-synaptic (**B**), and combined (**C**) NMJ morphological variables (*n* = 6). Correlation was assessed by calculating Pearson’s product moment correlation coefficient (*r*), the results of which are presented in **Supplementary Table 6** along with associated *P* values. No significant correlations were detected. Lumb. (foot), lumbricals of the foot; Lumb. (hand), lumbricals of the hand. See also **Supplementary Table 6**.

**Supplementary Figure 4.**
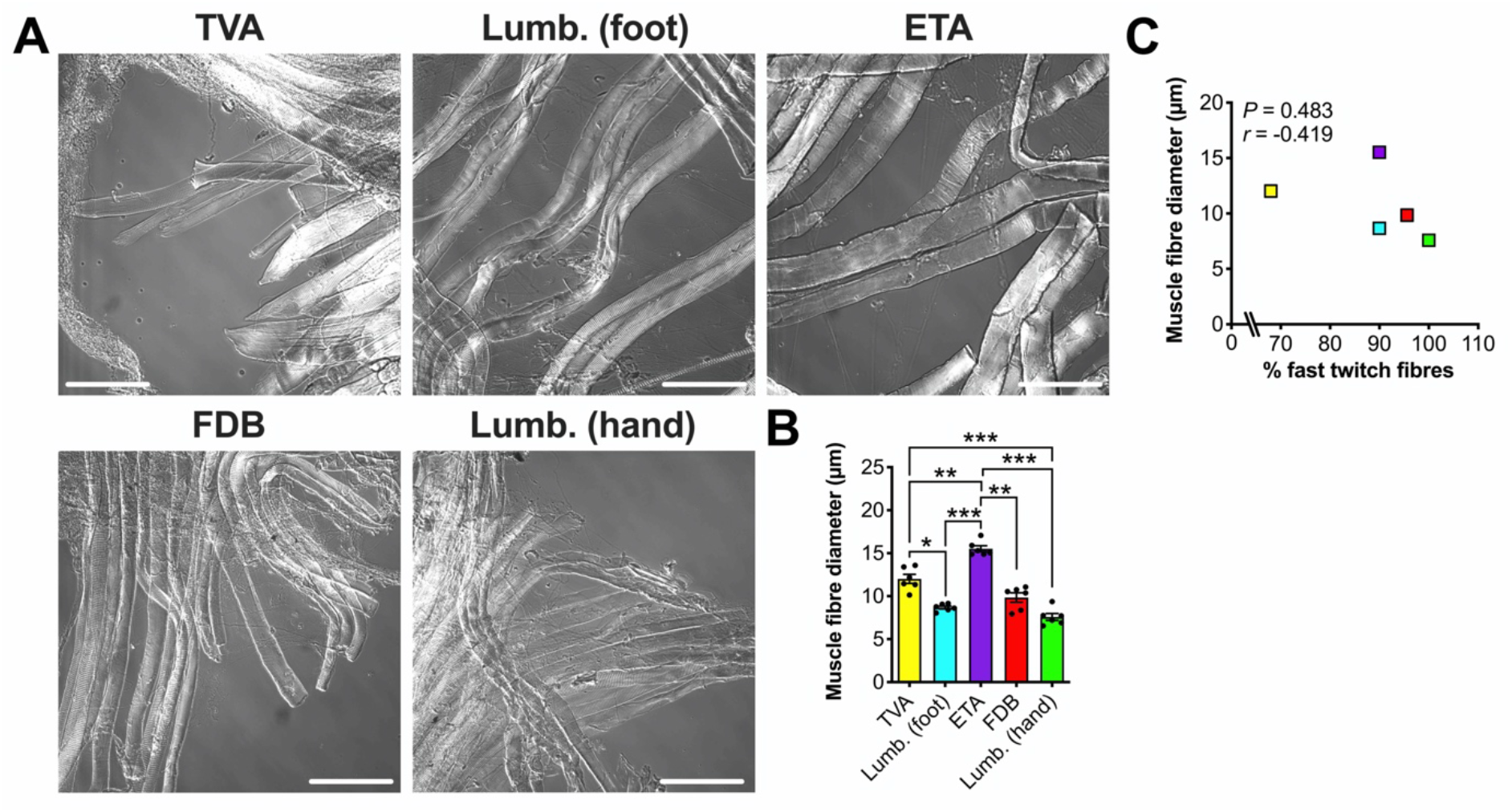
Muscle fibre diameters differ between muscles at P31-32, but do not correlate with fibre type. (**A**) Representative differential interference contrast images of teased muscle fibres from the five wholemount muscles. Scale bars = 100 μm. (**B**) Muscle fibre diameters are not equal across muscles. *P* < 0.001 repeated-measures one-way ANOVA. *P* < 0.05, **\*\****P* < 0.01, **\*\*\****P* < 0.001 Bonferroni’s multiple comparisons test. Means ± standard error of the mean are plotted (*n* = 6), as well as individual data points generated from each individual mouse (*i.e*. the mean values from 30 fibres). (**C**) There is no correlation between the percentage of fast twitch fibres found in each muscle (see **Table 1**) and fibre diameter. The colour-coding of muscles is maintained from panel B. *P* = 0.483, *r* = −0.419, Pearson’s product moment correlation. Lumb. (foot), lumbricals of the foot; Lumb. (hand), lumbricals of the hand.

**Supplementary Figure 5.**
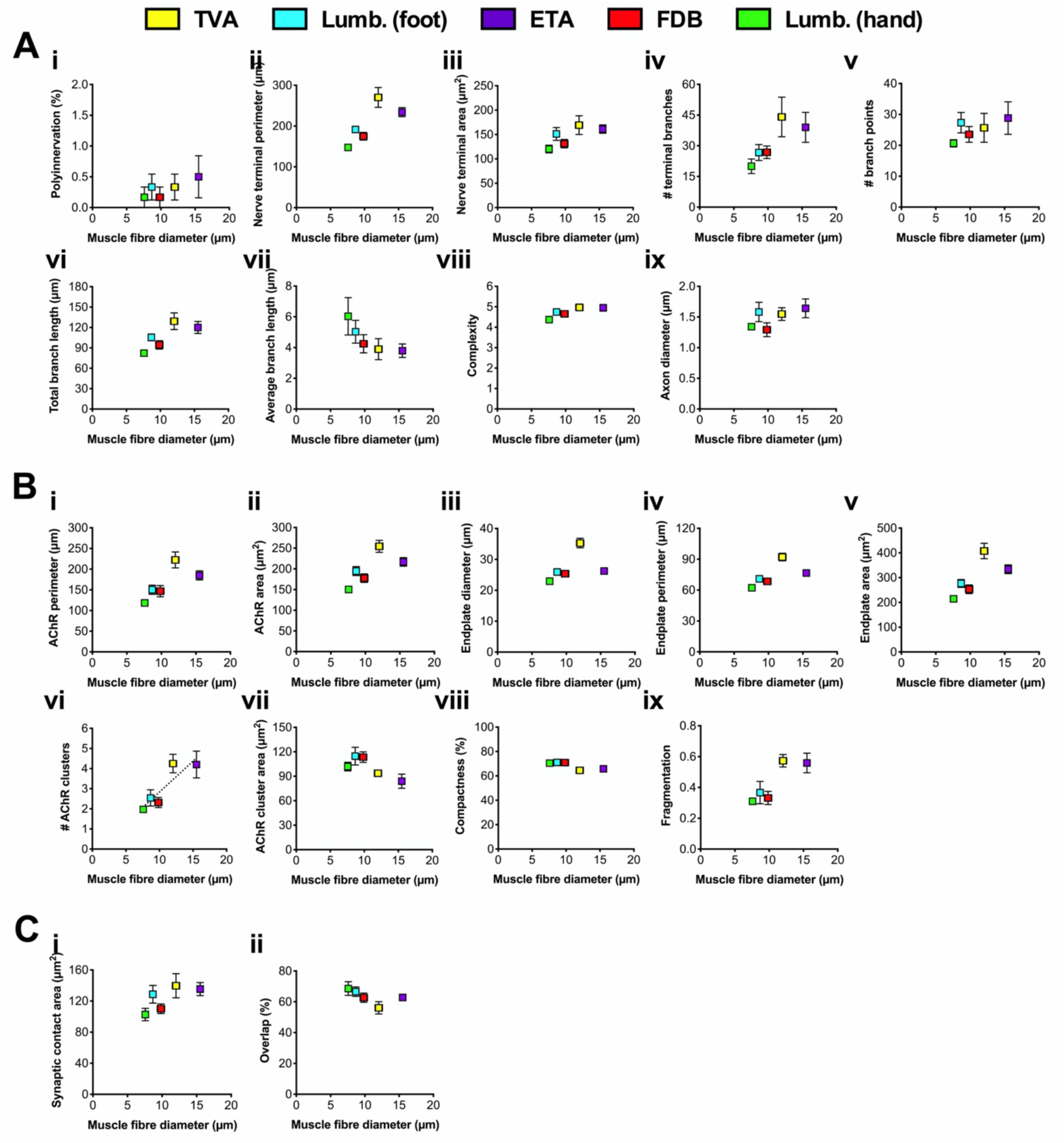
Muscle fibre diameter does not correlate with mature NMJ morphology. Muscle fibre diameter was correlated with all twenty pre-synaptic (**A**), post-synaptic (**B**), and combined (**C**) NMJ morphological variables from P31-32 muscles. Correlation was assessed by calculating Pearson’s product moment correlation coefficient (*r*), the results of which are presented in **Supplementary Table 7** along with associated *P* values. When the *P* value was < 0.05, a dashed line of best fit was plotted on the graph. Means ± standard error of the mean are plotted (*n* = 6). Lumb. (foot), lumbricals of the foot; Lumb. (hand), lumbricals of the hand. See also **Supplementary Table 7**.

**Supplementary Figure 6.**
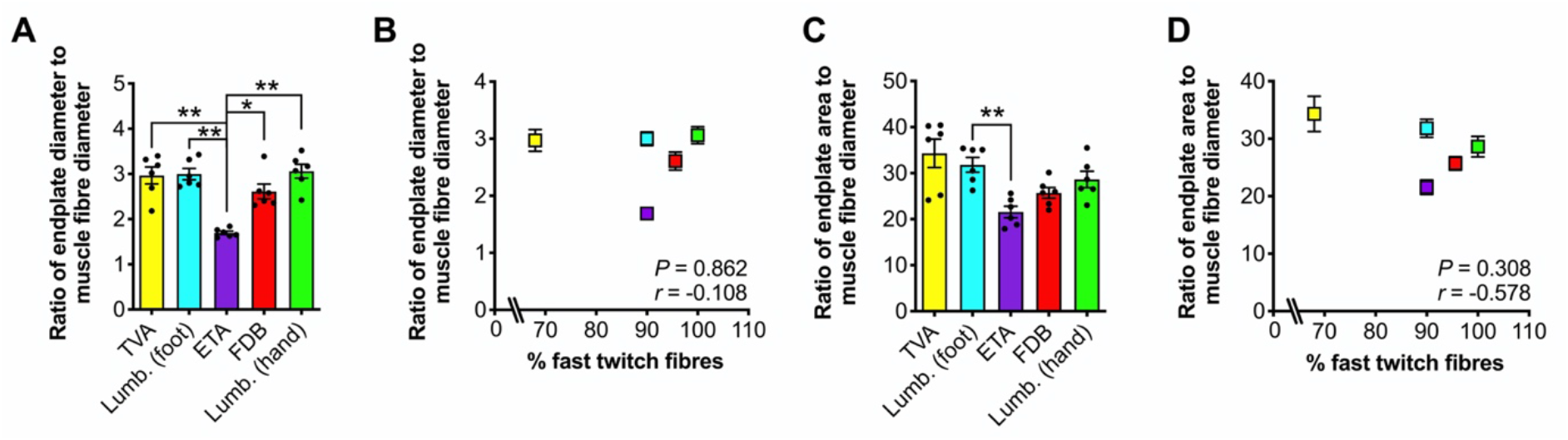
The ratios of endplate diameter and endplate area to muscle fibre diameter differ between muscles at P31-32, but do not correlate with fibre type. (**A**, **C**) The five wholemount muscles show different ratios of endplate diameter (A, *P* < 0.001) and endplate area (C, *P* = 0.002) to muscle fibre diameter (repeated-measures one-way ANOVA). *P* < 0.05, **\*\*** *P* < 0.01 Bonferroni’s multiple comparisons test. Means ± standard error of the mean are plotted (*n* = 6), as well as individual data points generated from each individual mouse. Lumb. (foot), lumbricals of the foot; Lumb. (hand), lumbricals of the hand. (**B, D**) There is no correlation between the percentage of fast twitch fibres found in each muscle (see **Table 1**) and the ratio of endplate diameter (B, *P* = 0.862, *r* = −0.108) or endplate area (D, *P* = 0.308, *r* = −0.578) to muscle fibre diameter (Pearson’s product moment correlation). The colour-coding of muscles is maintained from panels A and C.

## Supplementary Tables

**Supplementary Table 1.**
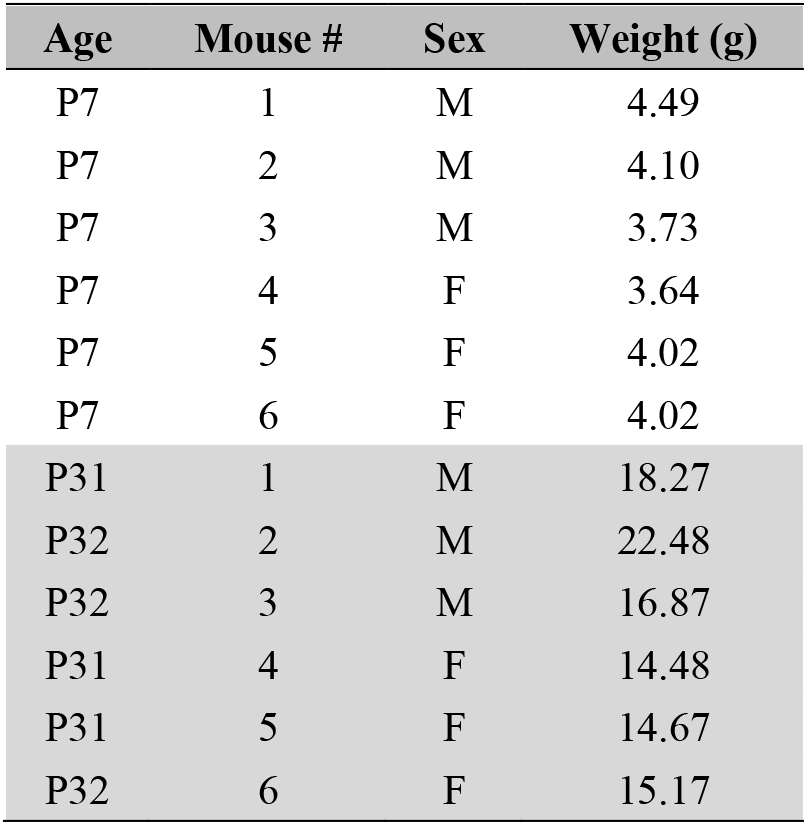
Details of mice used in this study. M, male; F, female.

| Age | Mouse # | Sex | Weight (g) |
| --- | --- | --- | --- |
| P7 | 1 | M | 4.49 |
| P7 | 2 | M | 4.10 |
| P7 | 3 | M | 3.73 |
| P7 | 4 | F | 3.64 |
| P7 | 5 | F | 4.02 |
| P7 | 6 | F | 4.02 |
| P31 | 1 | M | 18.27 |
| P32 | 2 | M | 22.48 |
| P32 | 3 | M | 16.87 |
| P31 | 4 | F | 14.48 |
| P31 | 5 | F | 14.67 |
| P32 | 6 | F | 15.17 |

**Supplementary Table 2.** Statistical testing of P31-32 NMJ morphological variables presented in Figure 2. Pre-synaptic variables are shaded green, post-synaptic variables shaded purple, and combined pre- and post-synaptic variables are unshaded. ^#^ *P* < 0.00256 repeated-measures one-way ANOVA, *i.e.* the Bonferroni correction-adjusted *P* value for an α of 0.05 when performing 20 associated tests. ns, not significant; **\*** *P* < 0.05, **\*\*** *P* < 0.01, **\*\*\*** *P* < 0.001 Bonferroni’s multiple comparisons test. When the ANOVA *P* > 0.05, Bonferroni’s multiple comparisons test was not performed (−). The bottom row shows the total number of morphological variables that were significantly different between each pair of muscles. Total sig., total significant. See also **Figure 2**.

| Variable | ANOVA<br>*P* value | TVA vs. Lumb. (foot) | TVA vs. ETA | TVA vs. FDB | TVA vs Lumb. (hand) | Lumb. (foot) vs. ETA | Lumb. (foot) vs. FDB | Lumb. (foot) vs Lumb. (hand) | ETA vs. FDB | ETA vs. Lumb. (hand) | FDB vs. Lumb. (hand) |
| --- | --- | --- | --- | --- | --- | --- | --- | --- | --- | --- | --- |
| Polyinnervation (%) | 0.696 | - | - | - | - | - | - | - | - | - | - |
| Nerve terminal perimeter (μm) | **0.003** | *ns* | *ns* | *ns* | *ns* | *ns* | *ns* | * | *ns* | * | *ns* |
| Nerve terminal area (μm<sup>2</sup>) | 0.079 | - | - | - | - | - | - | - | - | - | - |
| # terminal branches | 0.090 | - | - | - | - | - | - | - | - | - | - |
| # branch points | 0.387 | - | - | - | - | - | - | - | - | - | - |
| Total branch length (μm) | **0.015** | *ns* | *ns* | *ns* | *ns* | *ns* | *ns* | *ns* | *ns* | *ns* | *ns* |
| Average branch length (μm) | 0.250 | - | - | - | - | - | - | - | - | - | - |
| Complexity | 0.062 | - | - | - | - | - | - | - | - | - | - |
| Axon diameter (μm) | 0.120 | - | - | - | - | - | - | - | - | - | - |
| AChR perimeter (μm) | **<0.001<sup>#</sup>** | *ns* | *ns* | *ns* | * | *** | *ns* | * | *ns* | ** | *ns* |
| AChR area (μm<sup>2</sup>) | **<0.001<sup>#</sup>** | *ns* | *ns* | * | ** | *ns* | *ns* | *ns* | *ns* | * | *ns* |
| Endplate diameter (μm) | **<0.001<sup>#</sup>** | * | * | ** | ** | *ns* | *ns* | *ns* | *ns* | ** | *ns* |
| Endplate perimeter (μm) | **<0.001<sup>#</sup>** | * | *ns* | ** | ** | *ns* | *ns* | * | *ns* | ** | *ns* |
| Endplate area (μm<sup>2</sup>) | **<0.001<sup>#</sup>** | *ns* | *ns* | * | ** | * | *ns* | *ns* | *ns* | ** | *ns* |
| # AChR clusters | **0.003** | *ns* | *ns* | *ns* | * | *ns* | *ns* | *ns* | *ns* | *ns* | *ns* |
| AChR cluster area (μm<sup>2</sup>) | **0.046** | *ns* | *ns* | *ns* | *ns* | *ns* | *ns* | *ns* | *ns* | *ns* | *ns* |
| Compactness (%) | **0.004** | *ns* | *ns* | *ns* | *ns* | ** | *ns* | *ns* | *ns* | *ns* | *ns* |
| Fragmentation | **0.001<sup>#</sup>** | *ns* | *ns* | * | * | *ns* | *ns* | *ns* | *ns* | *ns* | *ns* |
| Synaptic contact area (μm<sup>2</sup>) | 0.117 | - | - | - | - | - | - | - | - | - | - |
| Overlap (%) | 0.141 | - | - | - | - | - | - | - | - | - | - |
| Total sig. |  | 2 | 1 | 5 | 7 | 3 | 0 | 3 | 1 | 6 | 0 |

**Supplementary Table 3.** Statistical testing of P7 NMJ morphological variables presented in Figure 3. Pre-synaptic variables are shaded green, post-synaptic variables shaded purple, and combined pre- and post-synaptic variables are unshaded. ^#^ *P* < 0.00256 repeated-measures one-way ANOVA, *i.e.* the Bonferroni correction-adjusted *P* value for an α of 0.05 when performing 20 associated tests. *ns*, not significant; **\****P* < 0.05, **\*\****P* < 0.01, **\*\*\****P* < 0.001 Bonferroni’s multiple comparisons test. When the ANOVA *P* > 0.05, Bonferroni’s multiple comparisons test was not performed (−). The bottom row shows the total number of morphological variables that were significantly different between each pair of muscles. Total sig., total significant. See also **Figure 3**.

| Variable | ANOVA<br>*P* value | TVA vs. Lumb. (foot) | TVA vs. ETA | TVA vs. FDB | TVA vs Lumb. (hand) | Lumb. (foot) vs. ETA | Lumb. (foot) vs. FDB | Lumb. (foot) vs Lumb. (hand) | ETA vs. FDB | ETA vs. Lumb. (hand) | FDB vs. Lumb. (hand) |
| --- | --- | --- | --- | --- | --- | --- | --- | --- | --- | --- | --- |
| Polyinnervation (%) | <0.001<sup>#</sup> | * | ** | * | * | * | *ns* | *ns* | * | *ns* | *ns* |
| Nerve terminal perimeter (μm) | <0.001<sup>#</sup> | ** | *ns* | ** | *ns* | ** | *ns* | *ns* | *** | *ns* | *ns* |
| Nerve terminal area (μm<sup>2</sup>) | <0.001<sup>#</sup> | ** | *ns* | ** | *ns* | *** | *ns* | *ns* | *** | * | * |
| # terminal branches | <0.001<sup>#</sup> | * | *ns* | *** | * | * | *ns* | *ns* | * | *ns* | *ns* |
| # branch points | <0.001<sup>#</sup> | * | *ns* | ** | * | * | *ns* | *ns* | ** | *ns* | *ns* |
| Total branch length (μm) | <0.001<sup>#</sup> | ** | *ns* | ** | * | ** | *ns* | *ns* | *** | *ns* | *ns* |
| Average branch length (μm) | 0.089 | - | - | - | - | - | - | - | - | - | - |
| Complexity | <0.001<sup>#</sup> | ** | *ns* | *** | ** | ** | *ns* | *ns* | ** | *ns* | *ns* |
| Axon diameter (μm) | 0.300 | - | - | - | - | - | - | - | - | - | - |
| AChR perimeter (μm) | 0.007 | * | *ns* | ** | *ns* | *ns* | *ns* | *ns* | * | *ns* | *ns* |
| AChR area (μm<sup>2</sup>) | <0.001<sup>#</sup> | ** | *ns* | *** | ** | ** | *ns* | *ns* | *** | ** | ** |
| Endplate diameter (μm) | <0.001<sup>#</sup> | ** | *ns* | ** | ** | ** | *ns* | *ns* | * | * | *ns* |
| Endplate perimeter (μm) | <0.001<sup>#</sup> | ** | *ns* | ** | ** | ** | *ns* | *ns* | ** | * | * |
| Endplate area (μm<sup>2</sup>) | <0.001<sup>#</sup> | ** | *ns* | *** | * | ** | *ns* | *ns* | *** | * | * |
| # AChR clusters | 0.057 | - | - | - | - | - | - | - | - | - | - |
| AChR cluster area (μm<sup>2</sup>) | <0.001<sup>#</sup> | *** | *ns* | *** | ** | *ns* | *ns* | *ns* | * | *ns* | *ns* |
| Compactness (%) | <0.001<sup>#</sup> | *** | *ns* | ** | *ns* | ** | *ns* | *ns* | * | *ns* | *ns* |
| Fragmentation | 0.029 | *ns* | *ns* | *ns* | *ns* | *ns* | *ns* | *ns* | *ns* | *ns* | *ns* |
| Synaptic contact area (μm<sup>2</sup>) | <0.001<sup>#</sup> | *** | *ns* | *** | * | *** | *ns* | *ns* | *** | * | * |
| Overlap (%) | 0.062 | - | - | - | - | - | - | - | - | - | - |
| Total sig. |  | 15 | 1 | 15 | 11 | 13 | 0 | 0 | 15 | 7 | 5 |

**Supplementary Table 4.** Statistical testing of P7 and P31-32 NMJ morphological variables presented in Supplementary Figure 2. Pre-synaptic variables are shaded green, post-synaptic variables shaded purple, and combined pre- and post-synaptic variables are unshaded. On the left, two-way ANOVA *P* values for differences between age, muscle and their interaction are presented. On the right, the mean values of each morphological variable at P31-32 are presented as a percentage of the mean of the same variable at P7 for all five muscles. The *P* values of Sidak’s multiple comparisons testing between P7 and P31-32 for each muscle are also presented. ^#^ *P* < 0.00256 two-way ANOVA interaction, *i.e.* the Bonferroni correction-adjusted *P* value for an α of 0.05 when performing 20 associated tests. *ns*, not significant; **\****P* < 0.05, **\*\****P* < 0.01, **\*\*\****P* < 0.001 Sidak’s multiple comparisons test. The bottom row shows the mean percentage change for each muscle in bold, and the total number of morphological variables that were significantly different between timepoints for each muscle. Total sig., total significant. See also **Supplementary Figure 2**.

| Variable | Two-way ANOVA *P* values |  |  | TVA |  | Lumb. (foot) |  | ETA |  | FDB |  | Lumb. (hand) |  |
| --- | --- | --- | --- | --- | --- | --- | --- | --- | --- | --- | --- | --- | --- |
|  | Age | Muscle | Interaction | P31-32 as<br>% of P7 | *P* value | P31-32 as<br>% of P7 | *P* value | P31-32 as<br>% of P7 | *P* value | P31-32 as<br>% of P7 | *P* value | P31-32 as<br>% of P7 | *P* value |
| Polyinnervation (%) | <0.001<sup>#</sup> | <0.001<sup>#</sup> | <0.001<sup>#</sup> | 0 | *** | 3 | *** | 4 | *** | 1 | *** | 2 | *** |
| Nerve terminal perimeter (μm) | <0.001<sup>#</sup> | <0.001<sup>#</sup> | 0.016 | 189 | *** | 278 | *** | 180 | *** | 331 | *** | 165 | ** |
| Nerve terminal area (μm<sup>2</sup>) | <0.001<sup>#</sup> | <0.001<sup>#</sup> | 0.198 | 324 | *** | 619 | *** | 341 | *** | 745 | *** | 340 | *** |
| # terminal branches | <0.001<sup>#</sup> | <0.001<sup>#</sup> | 0.013 | 57 | ** | 76 | *ns* | 50 | *** | 94 | *ns* | 56 | *ns* |
| # branch points | <0.001<sup>#</sup> | <0.001<sup>#</sup> | <0.001<sup>#</sup> | 45 | *** | 115 | *ns* | 58 | ** | 133 | *ns* | 73 | *ns* |
| Total branch length (μm) | <0.001<sup>#</sup> | <0.001<sup>#</sup> | 0.033 | 152 | *** | 271 | *** | 157 | *** | 328 | *** | 169 | ** |
| Average branch length (μm) | <0.001<sup>#</sup> | 0.129 | 0.401 | 300 | ** | 383 | *** | 332 | ** | 345 | ** | 389 | *** |
| Complexity | 0.520 | <0.001<sup>#</sup> | <0.001<sup>#</sup> | 92 | *ns* | 112 | *ns* | 94 | *ns* | 118 | ** | 97 | *ns* |
| Axon diameter (μm) | <0.001<sup>#</sup> | 0.153 | 0.265 | 177 | *** | 187 | *** | 199 | *** | 171 | *** | 148 | ** |
| AChR perimeter (μm) | <0.001<sup>#</sup> | <0.001<sup>#</sup> | 0.008 | 156 | *** | 153 | * | 135 | * | 180 | *** | 97 | *ns* |
| AChR area (μm<sup>2</sup>) | <0.001<sup>#</sup> | <0.001<sup>#</sup> | 0.001<sup>#</sup> | 254 | *** | 430 | *** | 221 | *** | 560 | *** | 259 | *** |
| Endplate diameter (μm) | <0.001<sup>#</sup> | <0.001<sup>#</sup> | 0.010 | 163 | *** | 200 | *** | 159 | *** | 206 | *** | 164 | *** |
| Endplate perimeter (μm) | <0.001<sup>#</sup> | <0.001<sup>#</sup> | 0.001<sup>#</sup> | 169 | *** | 196 | *** | 159 | *** | 207 | *** | 157 | *** |
| Endplate area (μm<sup>2</sup>) | <0.001<sup>#</sup> | <0.001<sup>#</sup> | <0.001<sup>#</sup> | 339 | *** | 444 | *** | 275 | *** | 546 | *** | 274 | *** |
| # AChR clusters | <0.001<sup>#</sup> | <0.001<sup>#</sup> | 0.007 | 104 | *ns* | 73 | *ns* | 94 | *ns* | 77 | *ns* | 43 | *** |
| AChR cluster area (μm<sup>2</sup>) | <0.001<sup>#</sup> | 0.527 | <0.001<sup>#</sup> | 242 | *** | 600 | *** | 250 | *** | 752 | *** | 500 | *** |
| Compactness (%) | <0.001<sup>#</sup> | 0.015 | <0.001<sup>#</sup> | 78 | *** | 97 | *ns* | 81 | *** | 103 | *ns* | 94 | *ns* |
| Fragmentation | <0.001<sup>#</sup> | <0.001<sup>#</sup> | 0.007 | 106 | *ns* | 74 | *ns* | 93 | *ns* | 71 | *ns* | 52 | *** |
| Synaptic contact area (μm<sup>2</sup>) | <0.001<sup>#</sup> | <0.001<sup>#</sup> | 0.255 | 329 | *** | 723 | *** | 336 | *** | 928 | *** | 389 | *** |
| Overlap (%) | <0.001<sup>#</sup> | 0.099 | 0.091 | 126 | * | 169 | *** | 147 | *** | 168 | *** | 148 | *** |
|  |  |  | Mean/Total sig. | 170 | 17 | 260 | 14 | 168 | 17 | 303 | 15 | 181 | 15 |

**Supplementary Table 5.** Statistical testing of correlation between the percentage of fast twitch muscle fibres and NMJ morphological variables at P31-32 and P7. Pre-synaptic variables are shaded green, post-synaptic variables shaded purple, and combined pre- and post-synaptic variables are unshaded. ^#^ *P* < 0.00256 Pearson’s product moment correlation, *i.e.* the Bonferroni correction-adjusted *P* value for an α of 0.05 when performing 20 associated tests. See also **Figure 4**.

|  | P31-32 |  | P7 |  |
| --- | --- | --- | --- | --- |
| Variable | *r* | Pearson<br>*P* value | *r* | Pearson<br>*P* value |
| Polyinnervation (%) | -0.409 | 0.494 | -0.771 | 0.127 |
| Nerve terminal perimeter (μm) | -0.912 | **0.031** | -0.698 | 0.190 |
| Nerve terminal area (μm<sup>2</sup>) | -0.841 | 0.075 | -0.652 | 0.233 |
| # terminal branches | -0.860 | 0.062 | -0.713 | 0.176 |
| # branch points | -0.410 | 0.494 | -0.783 | 0.117 |
| Total branch length (μm) | -0.871 | 0.055 | -0.726 | 0.165 |
| Average branch length (μm) | 0.603 | 0.282 | 0.288 | 0.639 |
| Complexity | -0.762 | 0.134 | -0.696 | 0.192 |
| Axon diameter (μm) | -0.516 | 0.373 | -0.198 | 0.750 |
| AChR perimeter (μm) | -0.929 | **0.022** | -0.548 | 0.339 |
| AChR area (μm<sup>2</sup>) | -0.939 | **0.018** | -0.664 | 0.222 |
| Endplate diameter (μm) | -0.992 | **<0.001<sup>#</sup>** | -0.898 | **0.038** |
| Endplate perimeter (μm) | -0.979 | **0.004** | -0.796 | 0.107 |
| Endplate area (μm<sup>2</sup>) | -0.942 | **0.017** | -0.632 | 0.253 |
| # AChR clusters | -0.770 | 0.128 | -0.064 | 0.918 |
| AChR cluster area (μm<sup>2</sup>) | 0.338 | 0.533 | -0.799 | 0.105 |
| Compactness (%) | 0.783 | 0.117 | -0.686 | 0.202 |
| Fragmentation | -0.782 | 0.118 | -0.008 | 0.990 |
| Synaptic contact area (μm<sup>2</sup>) | -0.806 | 0.100 | -0.643 | 0.242 |
| Overlap (%) | 0.884 | **0.047** | -0.202 | 0.744 |

**Supplementary Table 6.** Statistical testing of correlation between the percentage of fast twitch muscle fibres and the percentage change in NMJ morphological variables from P7 to P31-32. Pre-synaptic variables are shaded green, post-synaptic variables shaded purple, and combined pre- and post-synaptic variables are unshaded. See also **Supplementary Figure 3**.

| Variable | P7 to P31-32 |  |
| --- | --- | --- |
|  | *r* | Pearson *P* value |
| Polyinnervation (%) | 0.382 | 0.526 |
| Nerve terminal perimeter ($\mu\text{m}$) | 0.226 | 0.715 |
| Nerve terminal area ($\mu\text{m}^2$) | 0.362 | 0.550 |
| # terminal branches | 0.301 | 0.622 |
| # branch points | 0.569 | 0.317 |
| Total branch length ($\mu\text{m}$) | 0.398 | 0.507 |
| Average branch length ($\mu\text{m}$) | 0.802 | 0.103 |
| Complexity | 0.471 | 0.424 |
| Axon diameter ($\mu\text{m}$) | -0.345 | 0.569 |
| AChR perimeter ($\mu\text{m}$) | -0.357 | 0.555 |
| AChR area ($\mu\text{m}^2$) | 0.326 | 0.593 |
| Endplate diameter ($\mu\text{m}$) | 0.310 | 0.612 |
| Endplate perimeter ($\mu\text{m}$) | 0.140 | 0.823 |
| Endplate area ($\mu\text{m}^2$) | 0.130 | 0.835 |
| # AChR clusters | -0.809 | 0.097 |
| AChR cluster area ($\mu\text{m}^2$) | 0.629 | 0.256 |
| Compactness (%) | 0.714 | 0.175 |
| Fragmentation | -0.875 | 0.052 |
| Synaptic contact area ($\mu\text{m}^2$) | 0.399 | 0.506 |
| Overlap (%) | 0.699 | 0.189 |

**Supplementary Table 7.** Statistical testing of correlation between muscle fibre diameter and NMJ morphological variables at P31-32. Pre-synaptic variables are shaded green, post-synaptic variables shaded purple, and combined pre- and post-synaptic variables are unshaded. See also **Supplementary Figure 5**.

| P31-32 |  |  |
| --- | --- | --- |
| Variable | *r* | Pearson<br>*P* value |
| Polyinnervation (%) | 0.839 | 0.076 |
| Nerve terminal perimeter (μm) | 0.749 | 0.145 |
| Nerve terminal area (μm<sup>2</sup>) | 0.722 | 0.168 |
| # terminal branches | 0.823 | 0.087 |
| # branch points | 0.727 | 0.164 |
| Total branch length (μm) | 0.770 | 0.128 |
| Average branch length (μm) | -0.836 | 0.078 |
| Complexity | 0.822 | 0.088 |
| Axon diameter (μm) | 0.647 | 0.238 |
| AChR perimeter (μm) | 0.721 | 0.170 |
| AChR area (μm<sup>2</sup>) | 0.681 | 0.206 |
| Endplate diameter (μm) | 0.396 | 0.509 |
| Endplate perimeter (μm) | 0.592 | 0.293 |
| Endplate area (μm<sup>2</sup>) | 0.697 | 0.191 |
| # AChR clusters | 0.889 | **0.044** |
| AChR cluster area (μm<sup>2</sup>) | -0.807 | 0.099 |
| Compactness (%) | -0.804 | 0.101 |
| Fragmentation | 0.870 | 0.055 |
| Synaptic contact area (μm<sup>2</sup>) | 0.712 | 0.178 |
| Overlap (%) | -0.577 | 0.309 |

